# MOB rules: Antibiotic Exposure Reprograms Metabolism to Mobilize *Bacillus subtilis* in Competitive Interactions

**DOI:** 10.1101/2024.03.20.585991

**Authors:** Yongjin Liu, Sandra LaBonte, Courtney Brake, Carol LaFayette, Adam P. Rosebrock, Amy A. Caudy, Paul D. Straight

## Abstract

Antibiotics have dose-dependent effects on exposed bacteria. The medicinal use of antibiotics relies on their growth-inhibitory activities at sufficient concentrations. At subinhibitory concentrations, exposure effects vary widely among different antibiotics and bacteria. *Bacillus subtilis* responds to bacteriostatic translation inhibitors by mobilizing a population of cells (MOB-Mobilized *Bacillus*) to spread across a surface. How *B.* subtilis regulates the antibiotic-induced mobilization is not known. In this study, we used chloramphenicol to identify regulatory functions that *B. subtilis* requires to coordinate cell mobilization following subinhibitory exposure. We measured changes in gene expression and metabolism and mapped the results to a network of regulatory proteins that direct the mobile response. Our data reveal that several transcriptional regulators coordinately control the reprogramming of metabolism to support mobilization. The network regulates changes in glycolysis, nucleotide metabolism, and amino acid metabolism that are signature features of the mobilized population. Among the hundreds of genes with changing expression, we identified two, *pdhA* and *pucA*, where the magnitudes of their changes in expression, and in the abundance of associated metabolites, reveal hallmark metabolic features of the mobilized population. Using reporters of *pdhA* and *pucA* expression, we visualized the separation of major branches of metabolism in different regions of the mobilized population. Our results reveal a regulated response to chloramphenicol exposure that enables a population of bacteria in different metabolic states to mount a coordinated mobile response.

## Introduction

A major determinant of bacterial fitness includes how bacteria negotiate competition from diverse species they may encounter. To understand how bacterial communities of competitive species take shape and function in nature, it is essential to identify the mechanisms they use to interact (Galet et al., 2015; Harrison et al., 2008; Jones et al., 2017; Piewngam et al., 2018; Traxler et al., 2012). Model systems for interactions have revealed some unanticipated insights into natural interactions and bacterial population dynamics. Antibiotics promote bacterial fitness through inhibition of competitors. However, laboratory studies reveal subtle activities for antibiotics in the context of interspecies interactions (Davies, 2006; Fajardo and Martínez, 2008; Linares et al., 2006), functioning for example as environmental cues at subinhibitory concentrations (Bader et al., 2003; Goh et al., 2002; Herold et al., 2005; Linares et al., 2006; Tsui et al., 2004). Although the natural history of bacteria includes competition through antibiotic-mediated antagonism (Brien and Wright, 2011; Davies et al., 2006), the full spectrum of antibiotic impacts on communities in natural environments remains an open question (Brien and Wright, 2011; Davies, 2006; Fajardo and Martínez, 2008). Beyond growth inhibition, examples include measured changes in gene expression and dramatic changes in growth and metabolism, bacterial development, biofilm formation, and motility (Alves et al., 2018; Hernandez-Valdes et al., 2020; Hogan and Kolter, 2002; Jones et al., 2017; McCully et al., 2019; Morales et al., 2013; Onaka et al., 2011; Orazi and O’Toole, 2017; Shank et al., 2011; Stacy et al., 2014; Traxler et al., 2012). Also reported are instances of apparent counterattack provoked by a competitor (Basler et al., 2013; Hoefler et al., 2012; LeRoux et al., 2015; Mavridou et al., 2018; Stubbendieck and Straight, 2015).

We discovered that subinhibitory amounts of bacteriostatic translation inhibitors induce *Bacillus subtilis* colony expansion by sliding motility, which we call colony mobilization (Liu et al., 2018). Our initial observation was that *Streptomyces venezuelae* cultured on the same agar medium, induces mobilization of *B. subtilis* through production of chloramphenicol. We hypothesized that colony mobilization is an adaptive physiological function that enables a *B. subtilis* population to escape antibiosis before the antibiotic concentration becomes inhibitory. The mobilization under subinhibitory exposure provides an experimental system to dissect the regulatory functions that control the response to antibiotics from a competitor.

*B. subtilis* is an excellent model system to understand complex changes in growth of a bacterial population. A network of regulators control diverse physiological functions, including biofilm formation, sporulation, competence, and motility. Under overlapping control of different regulators, these functions modulate within a single population, forming a heterogeneous collection of cells in different physiological states (Dubnau and Losick, 2006; Eldar et al., 2009; Liu et al., 2015; Norman et al., 2013; van Gestel et al., 2015; Yannarell et al., 2023). For example, sporulation is the end point of a developmental cascade controlled principally by Spo0A (Burbulys et al., 1991). Spo0A also regulates functions leading to biofilm formation, which requires additional, more narrowly defined regulatory proteins such as SinR (Kearns et al., 2005; Shafikhani et al., 2002). Thus, cellular levels and activity of Spo0A influence many different functions needed to maintain fitness (Fujita et al., 2005; Kovács, 2016; Mirouze et al., 2011; Molle et al., 2003). An overarching view of the *B. subtilis* regulatory architecture reveals a network of genetic functions that confer multiple options to respond dynamically to shifting environments (Cao and Kuipers, 2018; Lopez et al., 2009; López and Kolter, 2010). The regulatory modules coupled with sporulation, competence, biofilm formation, swimming, and swarming motility have been extensively studied. Arguably the least is known about regulation of *B. subtilis* sliding motility (Grau et al., 2015; Kinsinger et al., 2005; Liu et al., 2018).

Sliding is a communal form of expansive surface growth that is independent of flagellar-mediated motility (Kearns, 2010; Kinsinger et al., 2003). Sliding requires mechanical elements that include extracellular polysaccharides, the protein BslA, and the lipopeptide surfactin (Grau et al., 2015; Hölscher and Kovács, 2017; Kearns, 2010; Kinsinger et al., 2003; van Gestel et al., 2015). Typically sliding occurs when cells are plated on specialized, semi-solid media (0.3% agarose or 0.7% agar), resulting in surface expansion instead of formation of a typical colony. Other requirements for sliding include purine and pyrimidine metabolic pathways (Kinsinger et al., 2005) and branched-chain fatty acids (Grau et al., 2015). However, it is not known how different environmental stimuli may influence the control of sliding motility. The regulatory proteins Spo0A and AbrB, as well as the regulatory kinases KinB and KinC, were shown to be important for sliding motility under specific nutrient conditions (Grau et al., 2015). It is unknown how *B. subtilis* responds to antibiotics that trigger sliding motility (Liu et al., 2018).

Chloramphenicol exposure at inhibitory concentration induces a (p)ppGpp-dependent stress response, which is required to maintain cell viability when translation is blocked (Yang et al., 2022). Using transcriptional and metabolic profiling, we find that subinhibitory chloramphenicol exposure has an opposite effect, resulting in elevated purine levels and regulated changes in metabolism during *B. subtilis* mobilization. We organized the data into networks of regulatory functions that reveal spatiotemporal separation of different metabolic states. The nutrition-sensing regulatory protein, CodY, is a central node in this regulatory network. Using time-lapse recordings of the expanding population, we observe dynamic patterning of carbon and nitrogen metabolism that supports mobilization of *B. subtilis*. We propose a model of population-wide, regulated coordination of metabolism that mobilizes *B. subtilis* upon exposure to antibiotics from competitors and thereby enhances competitive fitness.

## Results

### Exposure to translation inhibitors causes a transient colony expansion of *B. subtilis*

Previously, we found *S. venezuelae* induces *B. subtilis* colony mobilization following subinhibitory exposure to a selection of different translation inhibitors, including chloramphenicol (Liu et al., 2018). We initially sought to determine whether the response to chloramphenicol is a transient outcome of regulated changes in growth or whether it arises from emergence of a mutant population. If the response were transient, we predicted that the sliding population would revert to normal growth in the absence of chloramphenicol. We induced colony mobilization using subinhibitory chloramphenicol and subsequently transplanted cells from the sliding population to agar plates containing different concentrations of chloramphenicol (0 µM- control, 1 µM- stimulatory, and 16 µM- inhibitory) (**Figure 1A and 1B, Movie 1 and 2**). When transplanted to fresh media without chloramphenicol, *B. subtilis* ceased surface expansion. In contrast, on fresh media containing the stimulatory concentration of chloramphenicol (1 µM), the colony expansion continued, indicating that chloramphenicol exposure drives the mobilization, as opposed to a mutant phenotype of dysregulated growth. Growth was inhibited at the minimal inhibitory concentration (MIC) (16 µM), indicating that the population had not acquired resistance to chloramphenicol (**Figure 1C**). These results lead us to conclude that colony expansion is a transient characteristic stimulated by chloramphenicol exposure.

**Figure 1.**
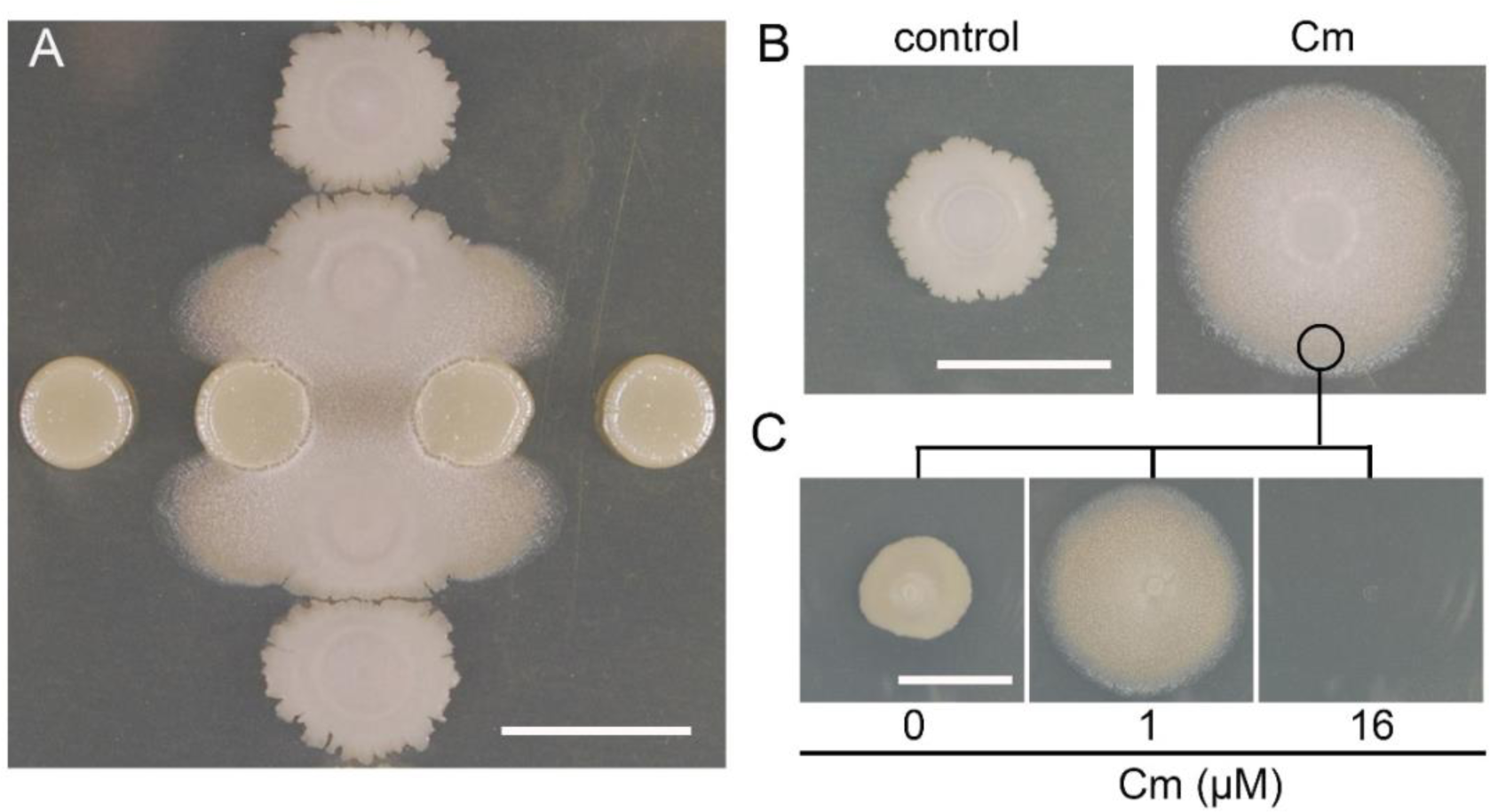
Subinhibitory chloramphenicol exposure drives *B. subtilis* mobilization. **A.** Sliding motility is induced in *B. subtilis* spots (vertical) when cocultured near *S. venezuelae* (horizontal spots) due to Cm exposure (imaged at 24 hours after inoculation, 30°C). **B.** Direct plating of *B. subtilis* on an agar plate with 1µM Cm leads to mobilization, and not in the control without Cm. **C.** Transplantation of *B. subtilis* cells from the mobilized population to different agar plates with the following concentrations of Cm: 0 μM, 1 μM (sub-MIC) and 16 μM (MIC). Bars, 1 cm.

We hypothesized that because different bacteriostatic translation inhibitors induced mobilization, and not antibiotics with other modes of action, suppression of protein synthesis may be the trigger (Liu et al., 2018). As a direct test we measured amino acid incorporation during protein synthesis in the presence of chloramphenicol, comparing chloramphenicol-treated to -untreated populations. Using the MIC (16 µM) for chloramphenicol, we observed a 60% reduction in amino acid incorporation following 5 minutes exposure (**Figure 2**). In contrast, when using the inducing concentration (1µM), we observed an approximately 7% reduction in amino acid incorporation relative to untreated controls over 5 minutes exposure. This modest reduction in amino acid incorporation under mobility-inducing conditions is consistent with the initiating event being a stress to translation that is insufficient to halt growth. Our observations support a model wherein regulated activation of colony mobilization is a response to translation stress, but the regulatory mechanism is undefined. To identify specific regulated functions associated with colony mobilization, we next considered how exposure to chloramphenicol affects gene expression under conditions where mobilization occurs.

**Figure 2.**
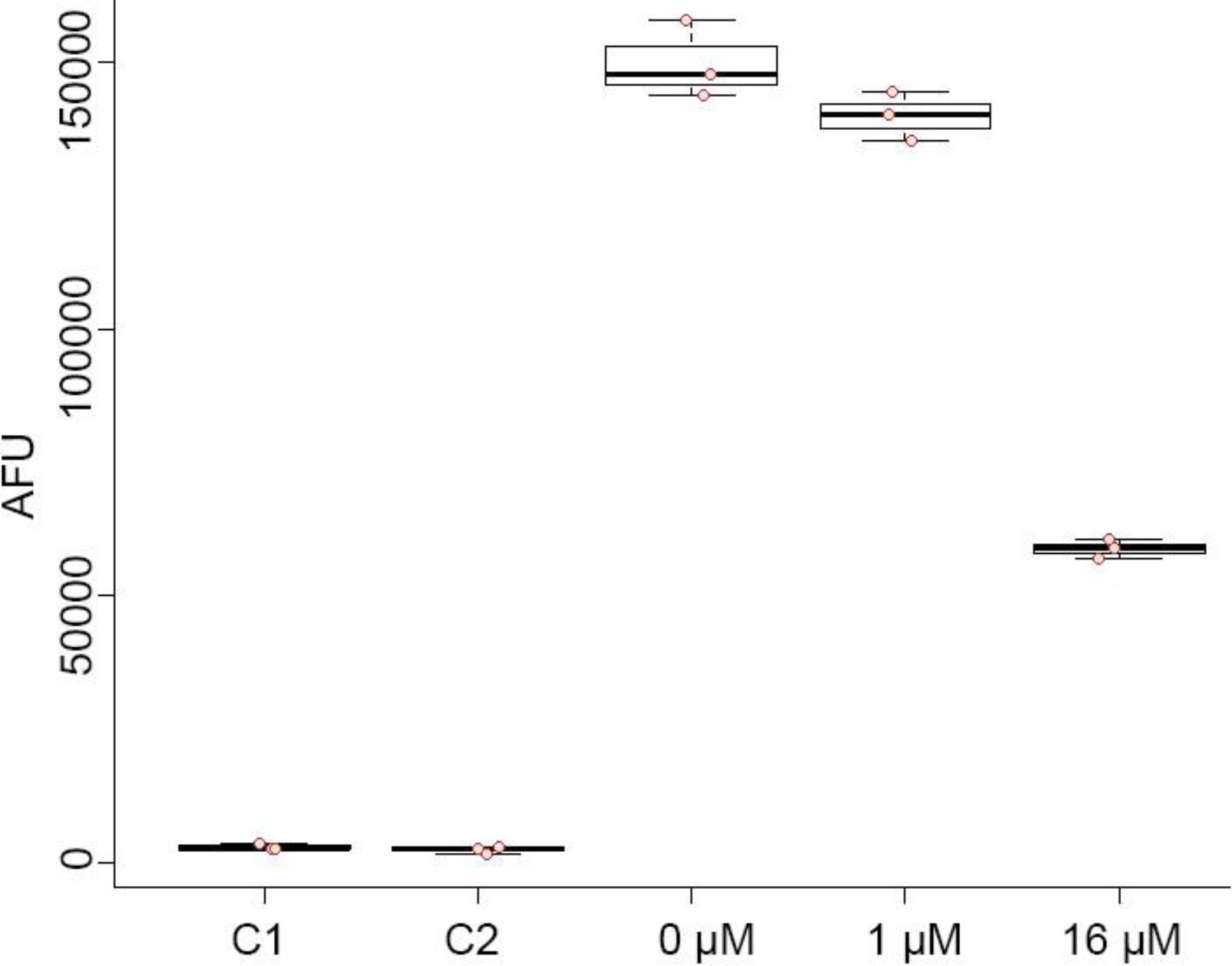
Subinhibitory chloramphenicol exposure supresses protein synthesis. The extent of protein synthesis was reduced in the presence of 1 μM subinhibitory concentration of Cm by Click-iT assay. C1 is a control without HPG. C2 is a control without Alexa Fluor 488. 16 µM Cm is equal to 1-fold MIC for Cm. The decrease of Cm-treated group was calculated based on the average of 0 µM Cm group. Each group contains three biological replicates (p value < 0.05 between 0 µM Cm and 1 µM Cm group). The cells were incubated with Cm for 5 minutes prior to sample fixation and processing.

### Patterns of gene expression suggest regulated changes in metabolism within the mobilized population

To identify changes in gene expression related to colony mobilization, we performed transcriptional analyses of the whole *B. subtilis* genome in the absence and presence of subinhibitory chloramphenicol. The experiment differs from prior studies by the concentration (1 µM) of chloramphenicol used and solid agar media instead of semi-solid media (Dar et al., 2016; Lin et al., 2005; Reilman et al., 2014). We focused on gene expression patterns at two times, using the reported expression pattern for *bmrCD* in response to chloramphenicol exposure as a guide (Lin et al., 2005; Liu et al., 2018; Reilman et al., 2014). Elevated *bmrCD* expression on agar media is transient with chloramphenicol, increasing from 4-6 hours and subsequently decreasing from 12-24 hours (Liu et al., 2018). We sampled cells based on the following criteria: 1) peak expression of the *bmrCD* operon as an indicator of early chloramphenicol exposure (6 hours), and 2) a later time point (24 hours) when colony expansion is visible. RNA sequencing results revealed that 793 and 352 genes are differentially expressed with chloramphenicol exposure, relative to untreated, at 6h and 24h, respectively (|fold change| ≥ 2 and adjusted p value <0.05).

We sorted the transcriptional data into functional categories using the assignments of genes in the Subtiwiki database (Zhu and Stülke, 2018). The resulting categorization displayed an enrichment of genes involved in several metabolic functions that exhibited a ≥ 2-fold change in expression with each condition (**Figure 3**). Within the functional categories identified, we observed multiple instances of expression patterns changing from majority increased to majority decreased or vice versa over the two times sampled. The observed temporal changes in gene expression suggested that the cells undergo a regulated transition from an initial response to chloramphenicol exposure (6 hours) into colony expansion (24 hours). This pattern was evident in different functional categories, including metabolism of cofactors, ribosomal biogenesis and translation, amino acid metabolism, and utilization of nitrogen source. For example, expression of ribosomal genes is transiently elevated at 6 hours, before colony expansion is visible, which may indicate an attempt by the cells to overcome chloramphenicol exposure by expanding the number of translating ribosomes and accelerating growth (Maaløe, 1979; Scott et al., 2010)(**Table S1**). Expression of genes involved in purine and pyrimidine metabolism were elevated at 6 hours and 24 hours, possibly reflecting changes in GTP, ATP, or other nucleotide pools or their related signaling metabolites (e.g. cyclic-di-GMP, (p)ppGpp). Aside from metabolic functions, transcriptional changes in other categories were observed. Genes in several functional categories are mostly repressed upon exposure to 1 µM chloramphenicol. For example, the general stress response genes (σB-regulon) are repressed, indicating that mobilization does not reflect a general stress response (Hecker et al., 2007; Hecker and Völker, 1998; McCormick et al., 2021; Price et al., 2001). To gain an organized, gene-specific view of the mobilization process, we focused on regulators that control genes within the identified categories.

**Figure 3.**
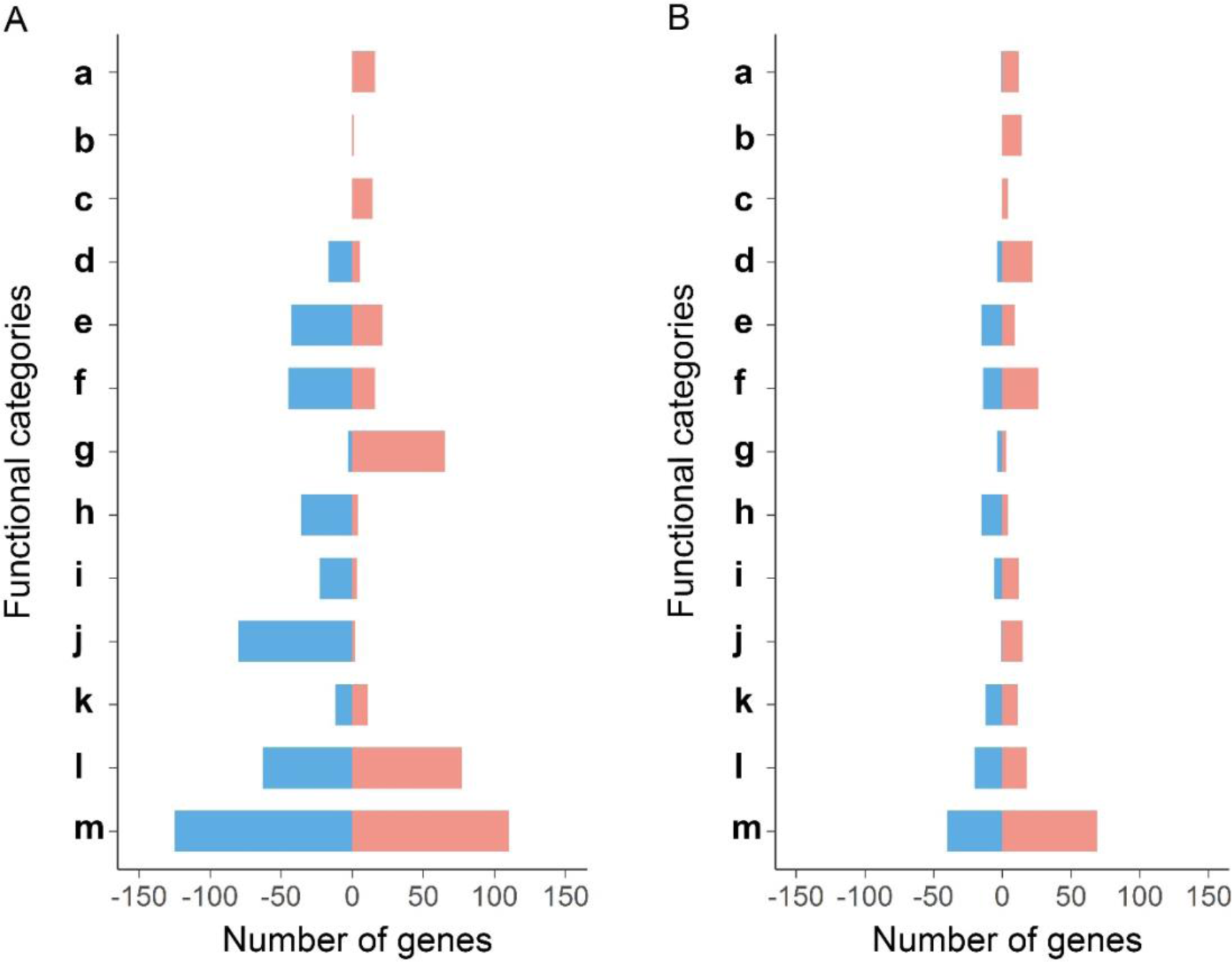
Patterns of gene expression reflect regulated changes in metabolism within the mobilized population. Functional classification of genes that change ≥2-fold with adjusted p value<0.05 in response to chloramphenicol at 6 h (**A**) and 24 h (**B**). **a.** Purine biosynthesis and salvage; **b.** Purine catabolism; **c.** Pyrimidine metabolism; **d.** Utilization of nitrogen sources; **e.** Carbon metabolism; **f.** Amino acid metabolism; **g.** Ribosomal biogenesis and translation; **h.** Secondary metabolism; **i.** Metabolism of cofactors; **j.** Stress response; **k.** Sporulation and germination; **l.** Others; m. unknown. Red: ≥ 2-fold increase, Blue: ≥ 2-fold decrease relative to control samples.

### A regulatory network view of chloramphenicol-induced colony surface expansion

To identify regulatory functions that may control colony mobilization, we used the *B. subtilis* regulons from SubtiWiki to sort genes that change expression under their respective regulators (Zhu and Stülke, 2018). We cataloged the identities and fold changes in gene expression for every regulon (**Table S2**). We then organized the outputs into networks that emerge at 6 and 24 hours, using the threshold of gene expression changing by ≥ 2-fold (**Figure 4A and 4B**). Our analysis revealed that for multiple regulators (e.g. AbrB, Abh, CodY, ResD) only a fraction of the genes in the regulon met the threshold for changing expression upon chloramphenicol exposure (Brinsmade et al., 2014)(**Table S2**). Therefore, we define the relative engagement of a regulon by the percentage of its genes that are expressed at least 2-fold up or down, relative to untreated controls. To visually highlight the roles of major regulators in response to chloramphenicol, we then applied a cutoff of 30% engagement for a regulator to appear in the network (**Figure 4**).

**Figure 4.**
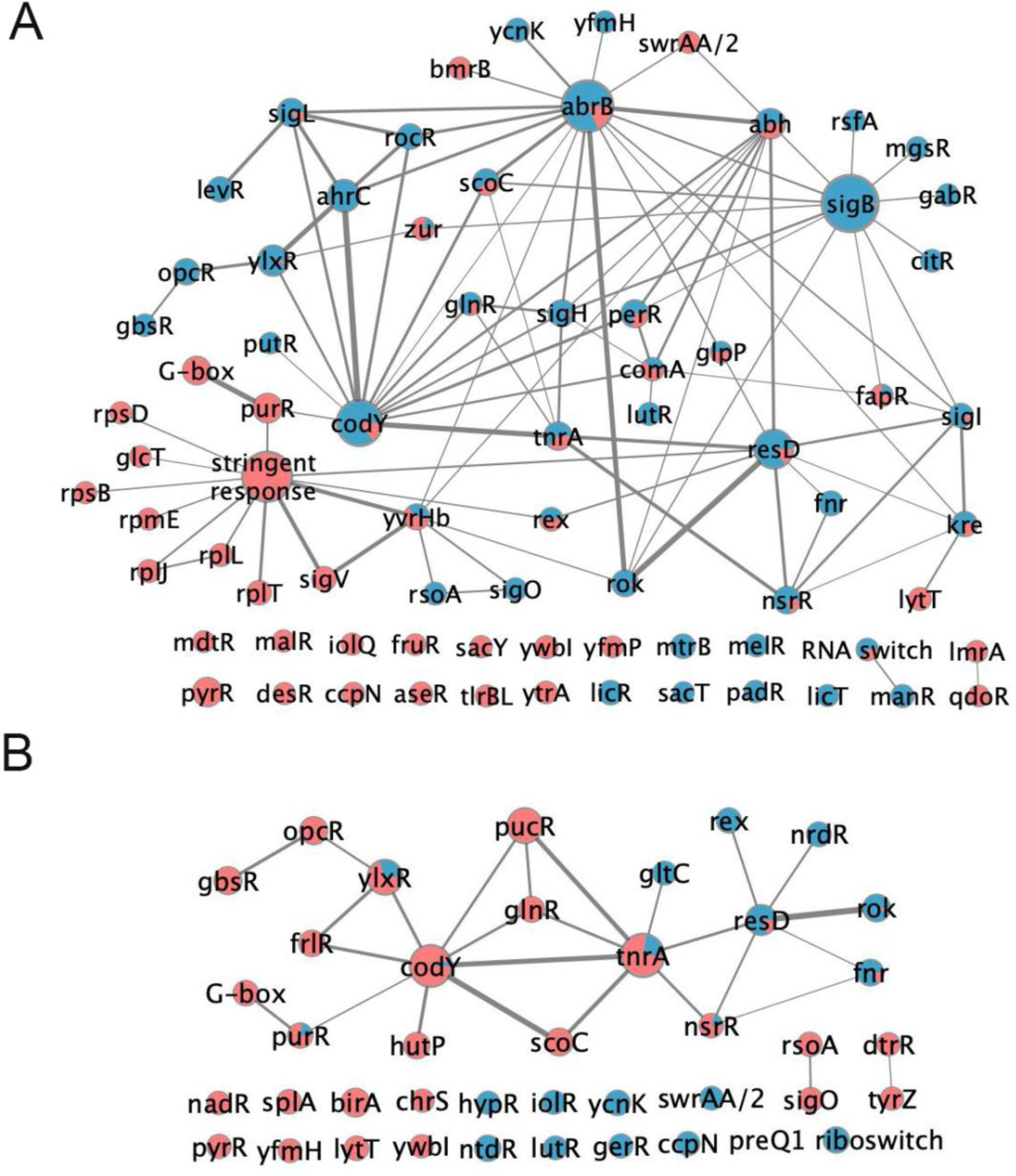
A network view of regulation highlights pathways of control for adaptive colony surface expansion. Cytoscape networks of regulators with ≥ 30% engagement of regulated genes at either: **A.** 6 h following chloramphenicol exposure (gene expression ≥ 2-fold difference from untreated controls with adjusted p-value < 0.05), or **B.** 24 h following chloramphenicol exposure (gene expression ≥ 2-fold difference from untreated controls with adjusted p-value < 0.05). In both A and B, the size of each node represents weighted gene number in each regulon (weighted by percentage of differentially expressed genes in each regulon) and the edge width represents weighted overlapped gene number between two regulons (weighted by percentage of differentially expressed genes among overlapped genes). Blue: downregulation; Red: upregulation.

We speculated that the engagement of different regulons under chloramphenicol exposure may reveal key features controlling colony mobilization. We identified major networked regulators based on two criteria: 1) majority activation of the regulon at either 6 or 24 hours, or 2) by a substantial switch between repressed and activated gene expression within the regulons from 6 to 24 hours. Under these criteria, prominent networked regulators include CodY, ScoC, PurR (G-box), GlnR, ResD, NsrR, LytT, FapR, TnrA, RnmP(YlxR), and PucR. The stringent response is engaged at 6 hours due to its control over ribosomal genes primarily. It is important to note that the observed pattern reveals overall suppression of the stringent response pathway, which is consistent with the elevated expression of purine biosynthetic genes and presumably elevated levels of GTP. In addition to networked regulators, we also identified network-independent regulators (no overlap between regulons) that met the 30% engagement threshold with chloramphenicol exposure (**Figure 4A and 4B**). The regulator identities emphasize major changes in metabolism in the mobilized *B. subtilis* population, including purine metabolism, nitrogen and carbon metabolism, and fatty acid metabolism.

### Identification of regulator mutants that disrupt *B. subtilis* mobilization

To determine how loss of regulator function impacts mobilization, we selected 9 of the major regulators (as classified above) for targeted gene deletions (**Figure 5**). We found for the networked regulators that only disruptions to CodY and PurR produced visible phenotypes on agar with or without subinhibitory chloramphenicol. Previous studies using soft-agar media showed that *B. subtilis* sliding motility depends upon two global regulatory proteins, AbrB and Spo0A (Grau et al., 2015). On the standard 1.5% agar media we used, a Δ*abrB* strain mobilized with or without 1 µM chloramphenicol, and the colonies had a rough biofilm-like appearance (**Figure 5**). The global regulator Spo0A is a repressor of AbrB that is implicated in regulation of sliding motility (Grau et al., 2015; Molle et al., 2003). Although Spo0A did not appear in the network (i.e. did not meet the engagement threshold), we tested a Δ*spo0A* strain for mobilization. A colony of the Δ*spo0A* strain produced a mucoid colony and did not mobilize on the agar media with or without chloramphenicol (**Figure S1**). The result is consistent with elevated AbrB repressor activity and no sliding motility in the absence of Spo0A. The major stress response regulator SigB appeared as a node in the 6-hour networks with most of the regulon repressed. We deleted the *sigB* gene to determine whether mobilization is impacted by loss of σB regulation (Binnie et al., 1986; Haldenwang and Losick, 1979; Hecker et al., 2007). No phenotype was observed for the Δ*sigB* strain, which is consistent with the reduced expression from the *sigB* operon observed in transcriptional profiles (**Table S1**).

**Figure 5.**
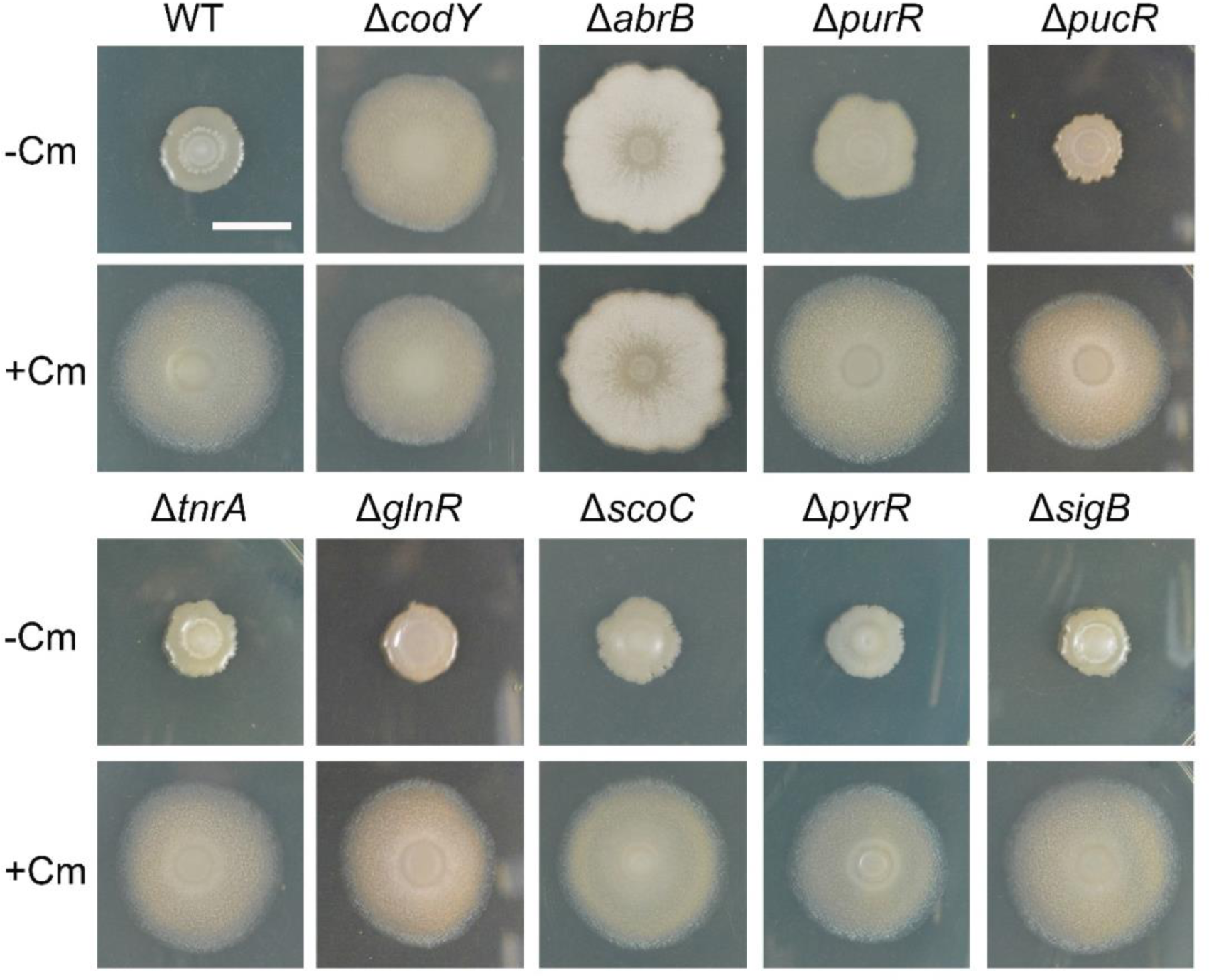
Genetic analysis of different regulators in the network orchestrating the sliding response. Strains with deletion of *codY*, *abrB*, *purR*, *pucR*, *tnrA*, *glnR*, *scoC*, *pyrR*, *sigB*, respectively, were analyzed in the absence (-Cm) and presence of Cm (+Cm). Pictures were taken at 24 h. Bar, 1 cm.

The Δ*codY, ΔabrB,* and Δ*purR* phenotypes in the absence of chloramphenicol suggest that within their regulons are genes that control sliding motility (**Figure 5**). Compared to AbrB and CodY, PurR controls a relatively small regulon (28 genes) required for the biosynthesis and metabolism of purines and contributes to other metabolic functions (e.g. purine salvage, formylation of Met-tRNA, isoleucine biosynthesis, and isoprenoid biosynthesis)(Saxild et al., 2001). The Δ*purR* strain exhibited sliding compared to wild type in the absence of chloramphenicol (**Figure 5**), although the extent of colony expansion was somewhat less when compared to the Δ*codY* strain. The Δ*purR* lesion results in depression of PurR-regulated genes, indicating a possible role for increase purine metabolites during mobilization. In addition to purine metabolism, pyrimidine biosynthesis has a reported role in sliding motility (Kinsinger et al., 2005; Turner et al., 1994). The PyrR-regulated pyrimidine biosynthesis regulon was upregulated with chloramphenicol exposure relative to the control (**Figure 4**). PyrR is a transcriptional attenuator that regulates pyrimidine biosynthetic genes, which are required for sliding motility (Kinsinger et al., 2005; Turner et al., 1998). Loss of *pyrR* function is expected to result in constitutive expression of the PyrR regulon (Turner et al., 1994). We used a deletion of the *pyrR* gene to determine if its loss of function, like *purR*, would lead to sliding motility without chloramphenicol, but the mutant strain did not produce a visible phenotype (**Figure 5**).

AbrB and CodY are master regulators with large regulons. AbrB regulates transcription of over 270 genes and controls the transition from growth to stationary phase (Banse et al., 2008; Chumsakul et al., 2013). Our transcriptomic data showed that the AbrB-engaged regulon was mostly repressed at 6 hours (101/272 genes showed ⩾2-fold change in expression, 18 up and 83 down) and did not reach the 30% engagement threshold for network inclusion at 24 hours (39/272 genes change expression ⩾ 2-fold, 16 up and 23 down). CodY is both an activator and repressor of genes in its large regulon (Brinsmade, 2016). CodY binds GTP and branched-chain amino acids (BCAAs) that are signals for nutrient abundance (Brinsmade et al., 2014; Sonenshein, 2005). Only a fraction, 76/188 and 65/188, of the CodY-regulated genes change expression ⩾ 2-fold at 6 and 24 hours, respectively. Among those genes at 6 hours, the majority are repressed, which is consistent with CodY being active and GTP and BCAAs levels sufficient to maintain its repressor activity (Brinsmade et al., 2014; Sonenshein, 2005). The direction of CodY-regulon activity reverses at 24 hours, when most of the engaged genes are derepressed. These differences suggest that the major regulators for nutrient and environmental sensing, CodY and AbrB (repressed by Spo0A), coordinately regulate the mobile response to chloramphenicol exposure. The network of their overlapping functions may define specific changes leading to chloramphenicol induced mobilization. AbrB and CodY share regulation of 34 genes. Of the shared genes, 25 showed differential expression ⩾2-fold following chloramphenicol exposure (**Table S2**). Three of the shared functions involve secondary metabolism (bacilysin and bacillaene) and an operon of unknown function (*yxbB-yxaM*), which includes an asparagine metabolic gene (*asnH*). A fourth shared function is for utilization of arginine, citrulline, and ornithine (*rocABC*). Although we have not determined whether specific shared genes are responsible for the constitutive mobilization observed with the Δ*abrB* and Δ*codY* strains, we suspect their complex phenotypes may arise from metabolic changes caused by dysregulation of their large regulons in the deletion strains.

CodY appears to play a key role in regulating mobilization due to its mutant phenotype and the engagement of its regulon at both 6 and 24 hours. We compared transcriptional changes in populations mobilized from chloramphenicol exposure versus a Δ*codY* strain to highlight differences between sliding motility induced by translational suppression versus the CodY regulator disruption. To determine the role of CodY in sliding motility, transcriptional analyses of the deletion strain was performed at 6 and 24 hours after plating on agar media identical to that used for chloramphenicol exposure. RNA sequencing results revealed that 1124 and 804 genes are differentially expressed in the deletion strain, relative to wildtype, at 6h and 24h respectively (|fold change| ≥ 2 and adjusted p value <0.05) (**Table S3**). As with the chloramphenicol-induced mobilization, the observed changes in gene expression were sorted into their relevant regulons (**Figure S4**). There are 101 and 74 regulators that meet the 30% threshold at 6 and 24 hours respectively, compared to 75 and 40 from chloramphenicol exposure. Comparing gene expression under chloramphenicol-induced versus Δ*codY*-constitutive mobilization, we identified early and late functional categories that appear specific to chloramphenicol-exposure based on the gene expression patterns (**Figure 6**). The early chloramphenicol-specific functions include purine biosynthesis (PurR), nitrate utilization (*nasBC*), and pyruvate uptake and utilization (LytT). At 24 hours, the chloramphenicol-specific gene expression patterns indicate sustained purine biosynthesis and nitrate utilization from 6 hours, and expand to include nitrite, purine, and histidine utilization (**Table S1**). In both *ΔcodY* and upon exposure to chloramphenicol at 6 hours, pyrimidine biosynthesis (PyrR), stringent response, ribosomal biosynthesis, and branched chain amino acid genes are upregulated (**Table S1** and **Table S3**) (**Figure 6**). Whereas at 24 hours, the shared gene expression patterns highlight pyruvate utilization, glutamine synthesis, NAD biosynthesis, and fructosamine utilization. Gene expression changes under chloramphenicol induction and constitutively mobilized Δ*codY* suggest that shared functions represent general mechanisms for sliding motility. Changes in gene expression specific to chloramphenicol induction at 6 hours, on the other hand, highlight possible triggers for mobilization and include elevated purine biosynthesis (PurR) and nitrate utilization (NadR).

**Figure 6.**
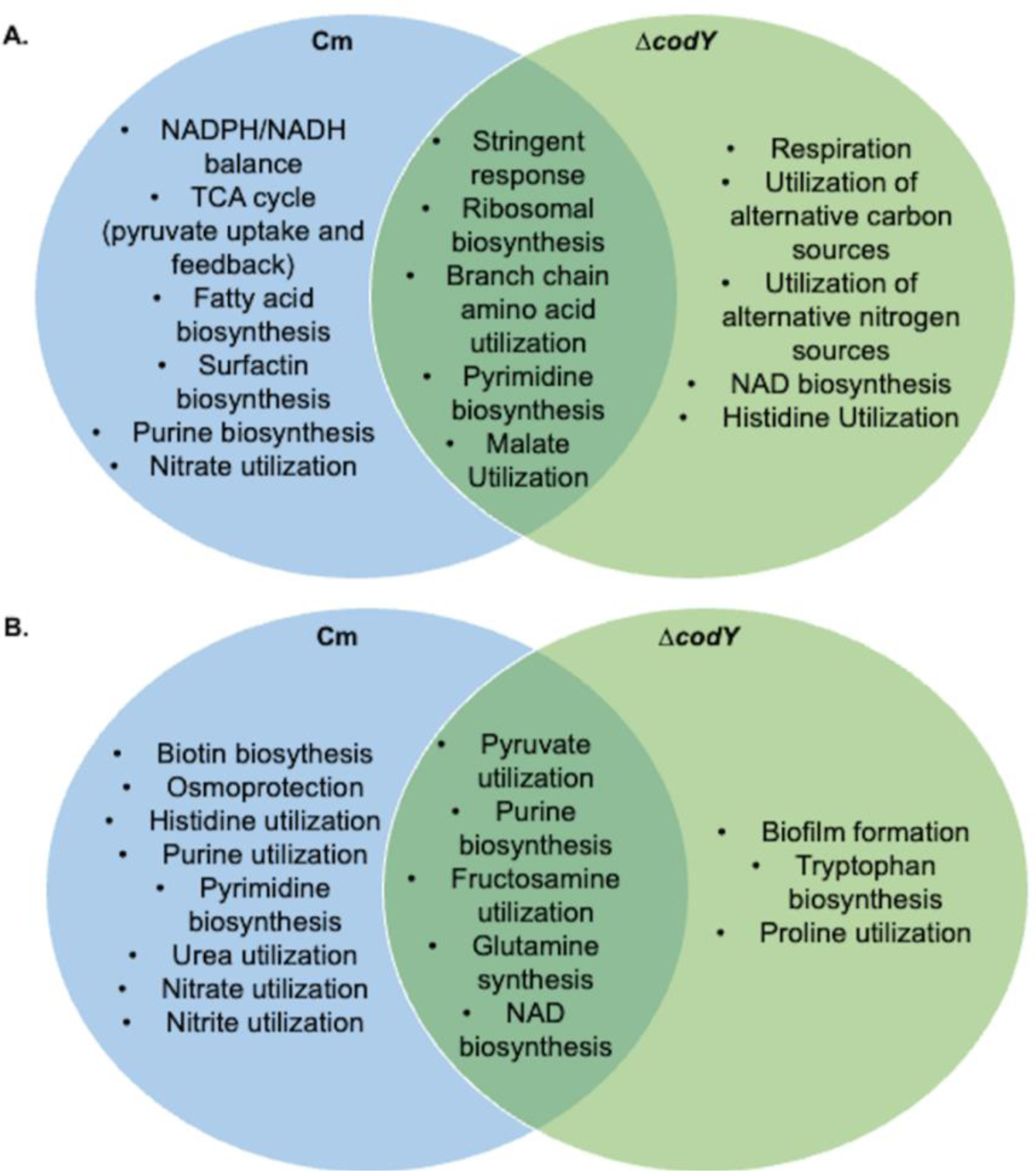
Comparison of major network regulators in wild type exposed to chloramphenicol and Δ*codY*. Venn diagram of prominent metabolic functions from regulon expression patterns in the Cytoscape networks for wild type exposed to chloramphenicol and Δ*codY* at **A.** 6 h or **B.** 24 h.

### Regulated changes in nucleotide metabolism for *B. subtilis* mobilization

The transcriptomic data indicates that purine metabolism has a role during mobilization, which is consistent with previously reported data (Kinsinger et al., 2005). PurR controls synthesis of purine metabolites (Weng et al., 1995). The PurR regulon is activated at 6 hours (between 2 and 20-fold among 17 out of 28 genes in PurR regulon), suggesting a drive for cells to synthesize purines following chloramphenicol exposure. The elevated expression of purine biosynthesis genes persists at 24 hours (between 2- and 8-fold among 9 out of 28 genes), although the fold activation and percent of genes activated diminished compared to 6 hours. Paradoxically, the majority of the PucR-regulated, purine catabolism regulon engages at 24 hours (5-468 fold) after chloramphenicol exposure, while purine biosynthetic genes remain upregulated (**Table S1**) (Beier et al., 2002; Schultz et al., 2001). However, the deletion strains for *pucR* as well as other regulator-encoding genes connected to *pucR* in the network, *tnrA, glnR,* and *scoC* did not show visible phenotypes for sliding motility in our assays (**Figure 5**). Thus, while purine biosynthesis promotes mobilization, purine catabolism appears dispensible for mobilization.

As direct test of a requirement for purine metabolism, we plated strains bearing mutations in *purD, purH,* and *pucM* with and without 1µM chloramphenicol (**Figure S2**). Although the *purD* and *purH* mutant strains require rich media for growth, they are competent to mobilize, revealing that the biosynthetic pathway itself is not essential for mobilization. The *pucM* deletion strain showed no visible phenotype relative to wild type in all conditions tested (**Figure S2).** In addition to purine metabolism, other apparent changes underscore a major reorganization of metabolism as cells mobilize. For instance, we observe elevated expression of both the histidine utilization genes and histidine biosynthetic genes at 24 hours (HutP and RnpM (YlxR) regulons, respectively) (**Table S2**). The differential regulation of histidine metabolism may connect purine and nitrogen metabolism through the production of AICAR (Rébora et al., 2005). The only gene co-regulated by CodY and PurR is *guaC*, which encodes a GMP-reductase that converts GMP to IMP through deamination of the purine base (Endo et al., 1983; Saxild et al., 2001). A deletion of the *guaC* gene did not have a visible phenotype for mobilization with or without chloramphenicol exposure (**Figure S3**). The transcription data indicate that chloramphenicol exposure leads to disruption of balanced metabolite pools, including purines and pyrimidines, but in concert with gene deletion phenotypes do not provide a clear candidate for a mobilization-triggering event. Because changes in transcription do not always reflect metabolic changes, we sought a direct measure of metabolite changes in the chloramphenicol-exposed cultures.

### Direct measurements of cellular metabolite pools indicate major changes in glycolysis and nucleotide metabolism during the transition to colony expansion

To determine how chloramphenicol exposure affects metabolism directly, we extracted metabolites from 6- and 24- hour populations of chloramphenicol-exposed, Δ*codY*, and control strains and quantitatively measured target metabolites by mass spectrometry. To illustrate the metabolomic profiles, we generated heatmaps displaying metabolites that either increased or decreased ≥ 1.5-fold at 6 hours (**Figure 7A, Figure S5A**), and ≥ 2-fold at 24 hours (**Figure 7B, Figure S5B**) (p-value < 0.05) compared to the levels measured in untreated controls (see **Table S4** for complete metabolite list). The resulting output from 6-hour samples revealed an increase in nucleoside triphosphates (NTPs) consistent with the observed transcriptional data (i.e. PurR and PyrR regulons) (**Figure 7A**). In addition to the elevated NTP levels, we observed diminished levels of precursor metabolites (NMPs, NDPs, and nucleosides) that are consistent with elevated flux in NTP biosynthesis following chloramphenicol exposure. Contrastingly, the Δ*codY* strain has increased levels of precursor metabolites (NMPs, NDPs, and nucleosides), in addition to NTPs. Other significant changes observed indicated enhanced activity in glycolysis through elevated levels of pyruvate and acetyl-CoA (**Table S4**) in both chloramphenicol-exposed and Δ*codY* cells (**Table S4**). Our metabolomic data reveal an increase in cellular aconitate, isocitrate, and citrate at 6 hours, with no changes detected in other TCA cycle metabolites (**Figure 7**). The changes in metabolite levels of greatest magnitude were consistent with increased demands for energy (pyruvate, phosphoenolpyruvate), nitrogen (glutamine), and metabolic enzyme activity (4-pyridoxic acid) (Richts et al., 2019), which we hypothesize generate a metabolic state under chloramphenicol exposure that leads to colony mobilization.

**Figure 7.**
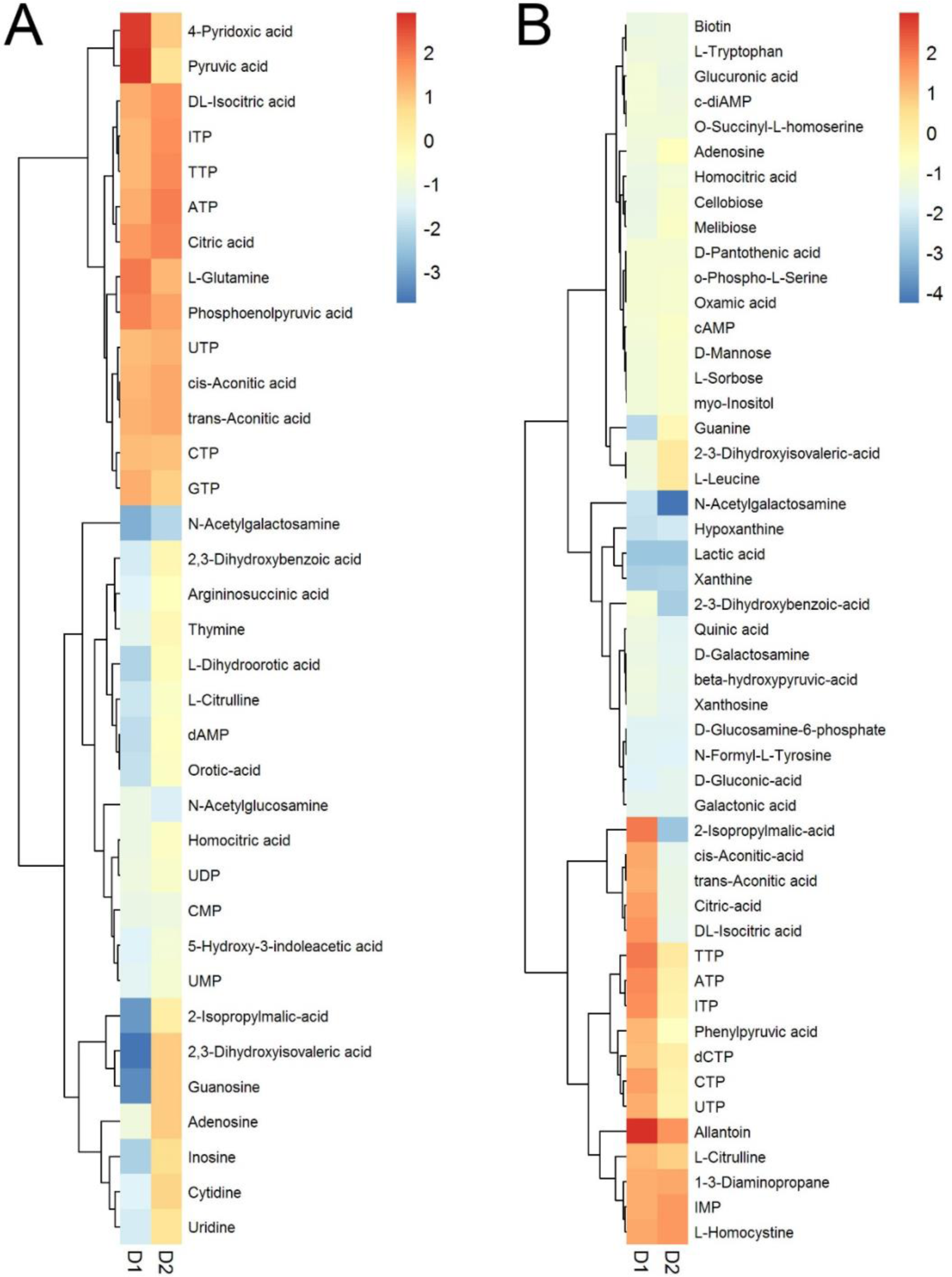
Metabolomics analysis underscores the pattern of shifting metabolism reflected by transcriptional analysis. Metabolic profiles of wild type at 6 h (**A**) and 24 h (**B**). Metabolites that change ≥1.5-fold (6 h) and ≥2-fold (24 h) upon exposure to 1 μM Cm in wild-type strain are listed (D1). The profiles of corresponding metabolites in Δ*codY* (D2) relative to wild-type strain at 6 h and 24 h are shown here for comparison.

Measurements at 24 hours revealed a larger set of metabolites with changing levels compared to untreated controls (**Figure 7B**). For the chloramphenicol-exposed cells, NTP levels remained elevated at 24 hours, consistent with measurements at 6 hours and the enhanced expression of PurR and PyrR regulons. Glycolysis and TCA cycle metabolites (pyruvate, citric acid, isocitrate) also remained elevated at 24 hours, indicating a sustained demand for carbon and energy to support colony mobilization. However, additional changes revealed elevated nitrogen metabolic activity, in particular purine catabolism (allantoin) and amino acid metabolism (arginine: L-citrulline, 1,3-diaminopropane, methionine: L-homocysteine, and phenylalanine: phenylpyruvic acid). Following early accumulation of NTPs in the cells, nitrogen may become a limiting factor in the mobilized population. By comparison, the *ΔcodY* cells lack the same metabolic demands observed in the chloramphenicol-exposed cells, for instance NTP levels and TCA cycle and glycolysis intermediates are reduced compared to the wild-type control.

The observed metabolic changes may provide clues to how the genetic regulatory networks control mobilization when exposed to chloramphenicol. Because CodY regulation is sensitive to both GTP and BCAAs, we considered the possibility that low levels of BCAAs may lead to activation of CodY-regulated genes. The branched-chain amino acid biosynthetic intermediates, 2-isopropylmalate and 2,3-diydroxyisovalerate, are reduced at 6 hours with chloramphenicol exposure relative to wild type, while they are elevated at 6 hours for the Δ*codY* strain (**Figure 7A**). Direct examination of BCAA levels however showed no significant reduction for isoleucine (1.1/1 -Cm-treated/-untreated) and leucine (0.99/1), and a modest decrease for valine (0.88/1) (**Table S4**). The observed BCAA levels are consistent with the majority of CodY-regulated genes being repressed at 6 hours given sufficient pools of BCAAs and GTP (Ratnayake-Lecamwasam et al., 2001; Shivers and Sonenshein, 2004). At 24 hours however, we observed a reduction in levels of both leucine (0.4/1) and valine (0.73/1) with chloramphenicol exposure (**Table S4**), compared to untreated controls, and no change in the Δ*codY* strain relative to wild type. No significant change was observed for isoleucine levels under any of the conditions tested. The observed changes in BCAAs and biosynthetic intermediates indicate a role for BCAA metabolism during sliding motility, but the changing levels may not directly influence CodY activity (Shivers and Sonenshein, 2004). This mode of CodY control is also supported by the observation that levels of GTP are elevated at 6 hours and not significantly different than controls at 24 hours (**Table S4**) (**Figure 7B**). An increased demand for BCAAs may arise from increased biosynthesis of BCFAs to support membrane biogenesis in the sliding population (Grau et al., 2015). To test whether BCAA levels have an impact on the colony expansion induced by Cm, we supplemented the medium with an excess of BCAAs (10mM each of isoleucine, leucine and valine) (Shivers and Sonenshein, 2004). However, this treatment did not substantially suppress or alter the chloramphenicol-induced sliding motility (**Figure S6**).

### A metabolic division of labor supports sliding motility in *B. subtilis*

We considered a model wherein chloramphenicol exposure creates a metabolic bottleneck that leads to reorganization of metabolism and a division of labor among cells in the population. The mobilized population reveals regulated changes in carbon and nitrogen use as the colony expands outward, including evidence of conflicting metabolic functions, such as elevated purine biosynthesis and catabolism within the population of cells, elevation of both histidine biosynthesis and utilization genes, and other similar patterns of oppositely directed metabolism (**Table S1**). We hypothesized that the cells may be dividing into different metabolic subpopulations that support mobilization. This form of division of labor among *B. subtilis* cells has been demonstrated in biofilm formation and motile populations (Chou et al., 2022; Liu et al., 2015; Norman et al., 2013; Rosenthal et al., 2018; van Gestel et al., 2015). A pattern of spatiotemporally separated metabolism was recently reported in a swarming population of cells (Jeckel et al., 2023).

We selected two metabolites that reflect major metabolic changes at 6 hours and at 24 hours as representatives of different metabolic states. At 6 hours with chloramphenicol, cellular levels of pyruvate (glycolysis, carbon metabolism) are elevated ≥3-fold over the untreated controls. Concurrently, transcripts of *pdhA* and *pdhB*, which encode pyruvate dehydrogenase, are 5-fold greater at 6 hours. Therefore, we used *pdhA* as a marker of early elevation of carbon metabolism upon chloramphenicol exposure. Allantoin is an intermediate in the catabolism of purines (Schultz et al., 2001). At 24 hours after chloramphenicol exposure cellular levels of allantoin (purine catabolism) are nearly 8-fold greater than controls (**Figure 7B**). Therefore, we focused on the *pucR* regulon to identify a marker of changes in nitrogen metabolism. We selected the *pucA-E* promoter based on the magnitude of the change in expression at 24 hours (≥ 267-468 fold over untreated controls).

To follow changes in gene expression in a colony of *B. subtilis*, we fused the promoter elements of *pdhA* and *pucA* to the luciferase reporter, *luxABCDE* (*lux*). Luciferase has the advantage of a relatively short signal turnover, which would enable us to resolve both increases and decreases in promoter activity (Brake et al., 2021; Popp et al., 2017; Radeck et al., 2013). We plated strains carrying the reporter fusions to agar media with or without chloramphenicol and monitored luciferase activity over time (**Figure 8, Movie 3**). The strains carrying the P*pdhA*-*lux* fusions revealed that the promoter is active at 6 hours in both treated and untreated samples. While the signal diminished over time in the untreated sample, it remained elevated in the chloramphenicol-exposed sample. Notably, at 48 hours we observed the peak of P*pdhA* activity on the periphery of the expanding colony, indicating that the cells with enhanced glycolytic activity are concentrated at the leading edge of the mobilized population. In contrast, the strains carrying the P*pucA*-*lux* reporter showed no detectable activity at early time points with or without chloramphenicol (**Figure 8, Movie 3**). Faint activity was observed beginning at 24 hours, which increased progressively thereafter. We observed that the P*pucA*-*lux* signal was internal to the expanding colony exposed to chloramphenicol, differentiating it from that of the peripheral localization of the P*pdhA*-*lux* signal. The results are consistent with a division of metabolic labor among cells in the expanding population, wherein elevated glycolysis defines the leading edge of the surface-expansive population, while compensatory changes in nitrogen metabolism may serve to rebalance metabolism within the subpopulation internal to the mobilized leading edge (Chou et al., 2022; Jeckel et al., 2023; Liu et al., 2015).

**Figure 8.**
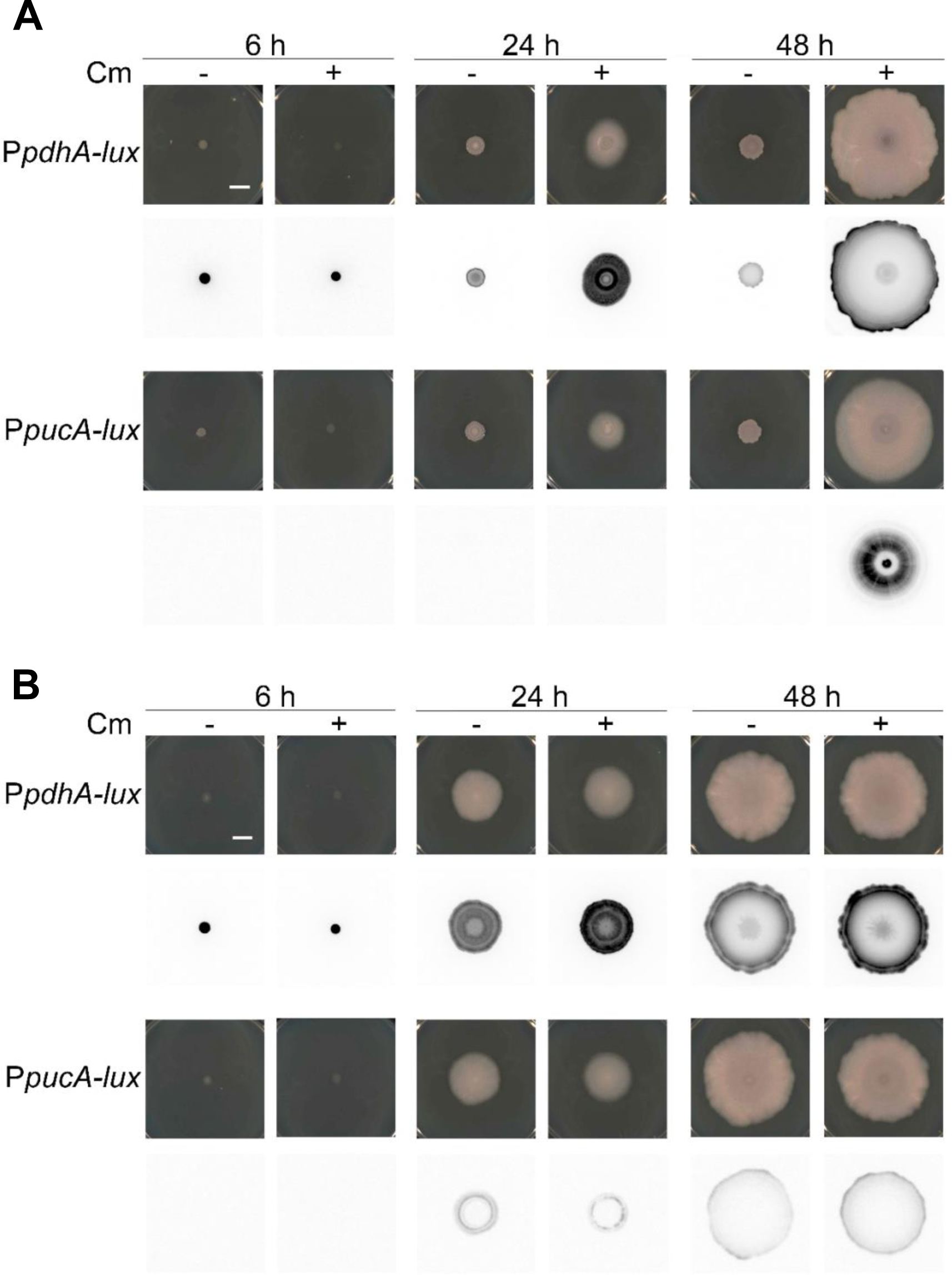
Spatial metabolism in the sliding population supports colony migration. Reporter strains with luciferase operon *luxABCDE* fused to *pdhA* or *pucA* promoter in the wild type *B. subtilis* (**A**) and Δ*codY* (**B**) strains were spotted on the agar plate without (-) or with Cm (+). Pictures were taken with phase contrast (top) and chemiluminescence (bottom) mode at different time points: 6 h, 24 h and 48 h. Bar, 1 cm.

We constructed the P*pdhA*-*lux* and P*pucA*-*lux* fusions in a Δ*codY* strain background. The strains were plated to media with and without chloramphenicol and sliding motility was observed in every case (**Figure 8B**). The P*pdhA*-*lux* expression pattern was now observed in both the chloramphenicol -treated and -untreated samples. Although the overall expression level is reduced compared to the population with chloramphenicol, elevated *pdhA* expression in Δ*codY* is consistent with elevated glycolytic activity as a driver of colony expansion. The P*pucA*-*lux* activity pattern was completely disrupted by the *codY* deletion. In these strains, we observed P*pucA*-*lux* activity at earlier time points and only faintly present at the colony edge, not the center as observed in wild type cells (**Figure 8B**). The observed changes indicate that, while chloramphenicol exposure enables *B. subtilis* to parse metabolic functions among subpopulations of mobilized cells, a defect in CodY regulation overrides spatiotemporal metabolic control to constitutively activate sliding motility.

Finally, we sought to identify phenotypes associated with mutations in glycolysis and purine catabolism. We spotted strains carrying deletions of *pdhA* (pyruvate dehydrogenase) and *pucM* (uricase: allantoin production) to agar plates with subinhibitory amounts of chloramphenicol (**Figure 9**). A deletion of *pdhA* disrupted sliding motility. Some changes were visible in colony morphology, even in the absence of sliding, compared to the wild-type strain. This result suggests that the cells respond to chloramphenicol exposure but cannot sustain the sliding population without elevated pyruvate dehydrogenase (glycolytic) activity. A deletion of *pucM* did not produce an apparent phenotype. Despite the substantial increase in expression of the *puc* genes and accumulation of allantoin, the expansion of the colony does not depend upon elevated purine catabolism.

**Figure 9.**
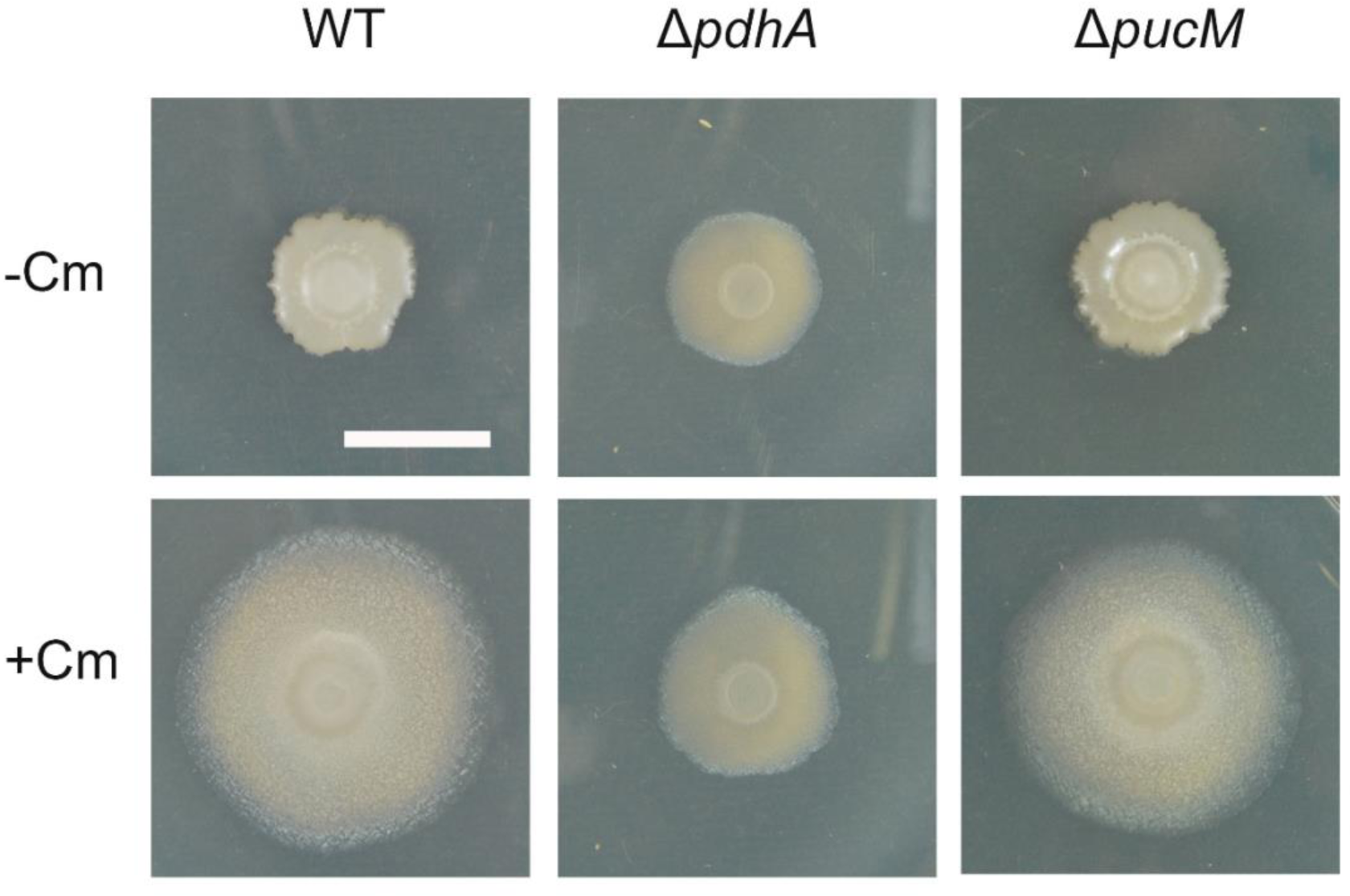
*pdhA* is required for colony expansion on the Cm plate. Wild type, Δ*pdhA, and* Δ*pucM* strains were spotted on the agar plate in the absence (-Cm) or presence of chloramphenicol (+Cm). Pictures were taken at 24 h. Bar, 1 cm.

**Figure 10.**
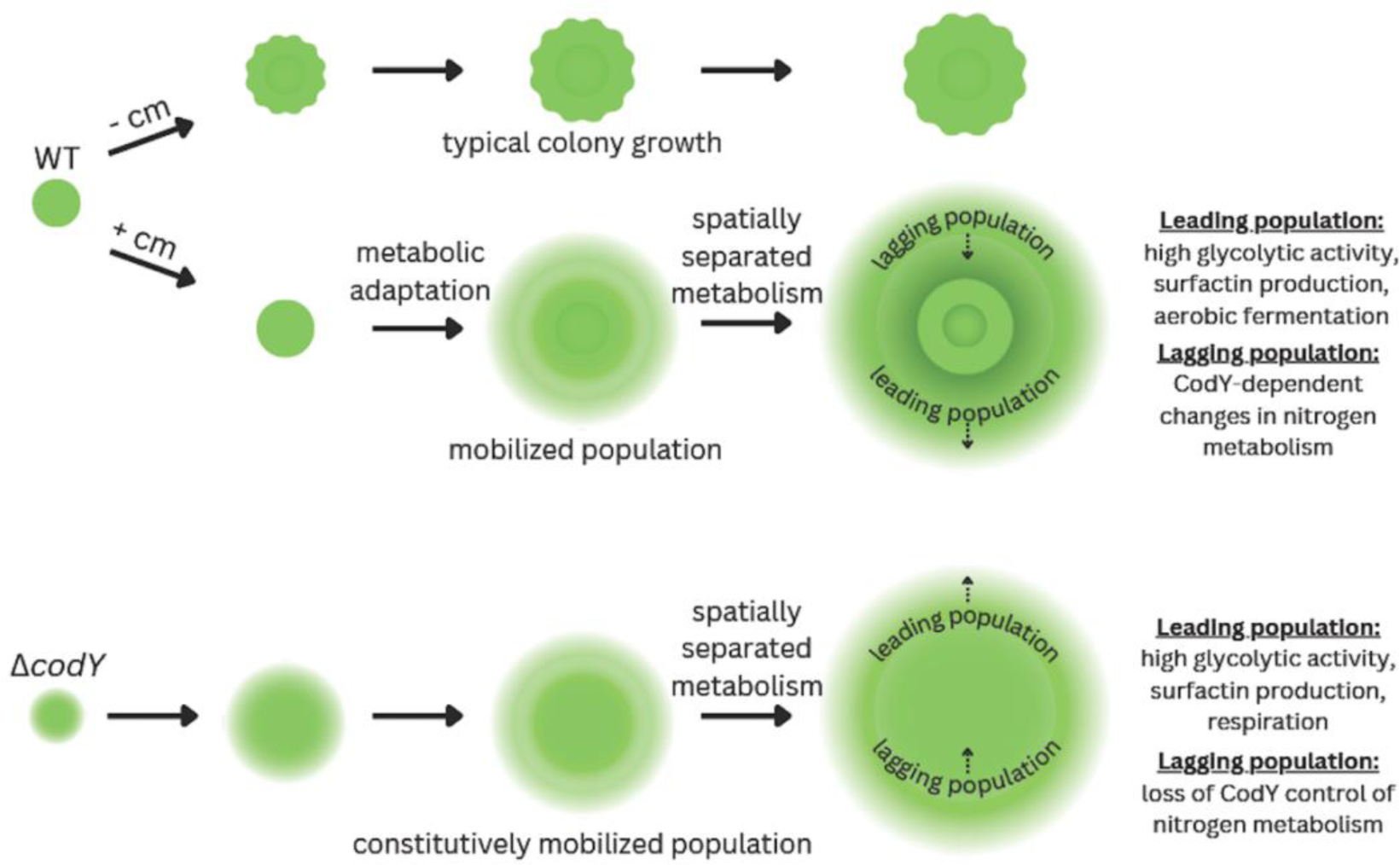
Model for chloramphenicol-induced mobilization using spatiotemporal control of metabolism compared to loss of CodY regulation. Wildtype *B. subtilis* growing on solid agar media develops a colony with typical size and morphology. Chloramphenicol exposure stresses translation, leading to a metabolic adaptation for colony expansion via sliding motility (mobilization). During outward growth of the population, spatiotemporal separation of different metabolic populations is visible using reporters for glycolysis and nitrogen metabolism respectively. A deletion of *codY* disrupts regulation of mobilization, resulting in a constitutively mobilized population. The elevation of glycolytic activity, which is required for mobilization, persists at the leading edge of the population. However, the Δ*codY* strain loses control of nitrogen metabolism in the lagging population, revealing that the changes in nitrogen metabolism are not essential for mobilizing the population.

## Discussion

### An adaptive physiological response to antibiotic-producing competitors

Our goal for this study was to gain insight into how *B. subtilis* mobilizes a colony in response to subinhibitory exposure to translation inhibitors. Using subinhibitory chloramphenicol as an inducer, our data reveal a modest reduction in protein synthesis in exposed cells, which suggests translation stress is a trigger for colony mobilization. Mapping changes in transcription to known regulons in *B. subtilis*, we identified a gene regulatory network that reveals dynamic metabolic coordination among cells in the mobilized population. Our analysis revealed a previously unknown role for CodY in regulation of sliding motility. Combining transcriptional and metabolomic data from chloramphenicol-exposed wild type and Δ*codY* strains, we identified a subset of gene-functional categories that are specific to chloramphenicol-induced mobilization of a *B subtilis* population (**Figure 6)**. We used targeted mutagenesis and metabolomics data with the regulatory networks to identify *pdhA* and *pucA-E* as signature metabolic genes to report the separation of major branches of metabolism among cells in the mobilized population. The data suggest that aerobic fermentation (glycolysis) supports growth-dependent sliding motility at the leading edge of the moving population (**Figure 9**). We observed that the leading edge is followed by a lagging population of cells in a different metabolic state. The changes we observe upon chloramphenicol exposure reveal a regulated, spatiotemporal division of metabolism.

### Suppressed translation promotes growth-dependent bacterial mobilization

Cellular metabolism is central to a growth-dependent mode of bacterial colony surface expansion. Our data supports a model where chloramphenicol exposure places a moderate stress on translation, mimicking a nutritional stress, despite the presence of abundant nutrients to support growth. Cells are attuned to changes in translation efficiency upon nutrient depletion (Bosdriesz et al., 2015; Davies et al., 2006; Groisman and Chan, 2021; Jacobson and Gillespie, 1968; McKinlay et al., 2020). The major regulator for *B. subtilis* nutrient sensing is CodY, which binds GTP and BCAAs as ligands that signal sufficient nutrient status (Ratnayake-Lecamwasam et al., 2001; Shivers and Sonenshein, 2004). A conserved cellular response to severe nutrient loss is the stringent response, wherein GTP is converted to (p)ppGpp, leading to reduced growth rate and dormancy (Hauryliuk et al., 2015; Kriel et al., 2014, 2012; Sonenshein, 2007). The stringent response protects *B. subtilis* from cell death, including chloramphenicol exposure. At an inhibitory concentration (12 µg/mL), chloramphenicol induces (p)ppGpp production in *B. subtilis,* protecting the cells from lethality (Yang et al., 2022). However, the *B. subtilis* response to subinhibitory amounts of chloramphenicol (∼0.3 µg/mL = 1µM) is paradoxical because the population enters a growth-dependent form of surface expansion upon moderate suppression of protein synthesis (∼7% reduced relative to control). Under this condition, elevated levels of GTP (and sufficient BCAAs) maintain repression for most CodY-regulated genes. However, our gene regulator network reveals subsets of genes that are differentially regulated by CodY and other regulators to control the mobilization response. The distinct outcomes associated with exposure to different levels of antibiotic underscore the importance of integrated regulatory networks that provide bacteria options for responding to external stresses and competition. In this model, the bacteria have an opportunity to escape antibiotic suppression of growth by adapting their metabolism to promote surface expansion.

### Metabolic coordination enables surface expansion of mobilized *B. subtilis*

*B. subtilis* colonies and biofilms are heterogeneous collections of cells in different physiological states (Lopez et al., 2009; Pisithkul et al., 2019; Rosenthal et al., 2018; van Gestel et al., 2015). Exposure to chloramphenicol, and presumably other protein synthesis inhibitors that induce mobilization (Liu et al., 2018), appears to structure the heterogenous population into a coordinated assembly of sub-populations that spread across a surface. Although a detailed mechanism connecting protein synthesis to the initiation of a mobilized response remains to be identified, the outcome of chloramphenicol exposure provides a window into the regulation of a complex metabolic transition in a population of bacteria. The regulatory networks we identified are snapshots of coordinated early (6 hours) and late (24 hours) changes in expression for hundreds of genes in the dynamic population. The transcriptional reporters used in this study, P*pdhA* and P*pucA* fused with luciferase, enable us to view the dynamic changes in metabolic regulation during colony expansion. We suggest that overlapping regulatory functions in a metabolically variegated population of *B. subtilis* provides precision controls for specialized groups of genes that control a complex response to competition. The observed patterns of regulated gene expression suggest that the bacterial population reorganizes cells in heterogeneous states into to a structured, functional output (mobilization) for a competitive advantage (Rosenthal et al., 2018).

The major metabolic rewiring induced by chloramphenicol emphasizes five predominant categories: glycolysis and fermentation, respiration-nitrate utilization, purine and pyrimidine biosynthesis, nitrogen metabolism, and branched-chain amino acid and fatty acid biosynthesis. A major metabolic shift occurs between early and late stages, where what appears to be a stress response at 6 hours transitions to a mobilized population by 24 hours. Consistent with a previous report of genetic requirements for *B. subtilis* sliding motility (Kinsinger et al., 2005), we see elevated levels of gene expression of the purine and pyrimidine biosynthesis genes and elevated levels of NTPs in the early stages. The changes in NTPs, along with an initial increase in ribosomal subunit gene expression and an elevated glycolytic activity suggests energy is initially directed toward overcoming translation stress. The same effect is not observed in the Δ*codY* strain, wherein the loss of CodY regulation promotes a constitutively mobile population. To adapt to chloramphenicol exposure, elevated gene expression of *pdhA*, *pckA*, *maeA*, and *maeN* likely increase glycolytic activity for colony growth. In concert, we observed elevated phosphoenolpyruvate (PEP) and pyruvate in chloramphenicol-exposed cells. The observed changes in glycolytic activity are consistent with a recent report of spatiotemporally separated metabolism in swarming population (Jeckel et al., 2023). The mobilized cells also activate pyruvate transport (*pftA* and *pftB*) along with acetate production (*ackA*), presumably to support cell growth and colony expansion through fermentative ATP synthesis in the absence of respiration (Grundy et al., 1993; Wolfe, 2005). The use of aerobic fermentation by the chloramphenicol-exposed population is further supported by the increase in assimilatory nitrate reductase genes (*nasBC*) and NADH dehydrogenase (*ndh*) (**Table S1**). In contrast, the *ΔcodY* strain appears to use cytochrome BC as terminal electron acceptor based on elevated expression of *cydABCD* in the mobile population (Winstedt and von Wachenfeldt, 2000). This contrast between chloramphenicol-exposed and Δ*codY* strains indicates that different mechanisms balance NADH + H^+^/NAD^+^ and energy levels, according to their differential regulatory status (Gyan et al., 2006). Other notable changes in metabolism include increased levels of branched-chain amino acids uptake and utilization genes present under both conditions (*braB*, *bkdB*, and *bkdAB*). This observation agrees with a previously reported need for branched-chain fatty acid biosynthesis during sliding, presumably to support membrane growth for the expanding population (Grau et al., 2015).

While at 6 hours our data indicates translation stress and metabolic adaptation, at 24 hours we observe visible features of the spatiotemporal division of labor in the mobilized population (**Figure 8**). Previous studies report surfactin and EPS production along with BslA and TasA when focused closely on the leading edge of the colony. These functions operate cooperatively, providing the mechanical requirements to reduce surface tension and enable outward-directed growth (Grau et al., 2015; van Gestel et al., 2015). Focusing on the entire population, our results show the leading edge to be a region of high glycolytic activity, likely providing the energy and substrates to support the mechanical requirements for growth-dependent surface expansion. This pattern is shared between sliding and swarming populations (Jeckel et al., 2023), the latter requiring surfactin production and flagella to promote motility in semi-solid media (Julkowska et al., 2005; Kearns and Losick, 2004). Perhaps the swarming population cooperates with a sliding sub-population to enable outward expansion under permissive conditions. Behind the leading edge of the chloramphenicol-induced mobile population, we observe cells in distinctly different metabolic states. Gene expression patterns at 24 hours emphasize changes in nitrogen metabolism (urease, purine catabolism, histidine biosynthesis and utilization). The use of the P*pucA*-luciferase reporter and time-lapse imaging indicate the inner subpopulation is in a state of dynamically changing gene expression and metabolism (**Figure 8 and Movie 3**). Deletions in regulatory genes (**Figure 5**), the *pucM* gene, and a different pattern of P*pucA-*luciferase expression in a Δ*codY* strain, reveal that these changes are not essential for sliding motility. However, our data suggests different metabolic solutions are available to the mobile populations depending on the initiating condition, nutrition dysregulation versus translation stress. We hypothesize that the metabolic changes driving the leading edge of the mobile populations (chloramphenicol-induced versus Δ*codY*) are differentially compensated in the overall population depending upon available regulatory possibilities. For example, apparent futile cycling of histidine synthesis and degradation could provide a mechanism to supply AICAR for purine synthesis that is only available to the chloramphenicol-induced population, which have CodY function intact (Rébora et al., 2005). An important contributor to the metabolically available functions could be the residual nutrients following passage of the leading edge.

*Bacillus subtilis* evolved intricate regulatory systems to control cell fates in response to diverse competitors and extracellular cues. The chloramphenicol-mobilized *B. subtilis* reveals a population that reorganizes metabolism to support an adaptive physiological response to antibiotic exposure. An emphasis on fermentation and nitrogen regulation resembles the Crabtree effect (Warburg effect in cancer cells), where products of glycolysis are fed to biosynthetic pathways that support rapid growth (Erez and Kolodkin-Gal, 2017; Kukurugya and Titov, 2023; McKinlay et al., 2020). Our data suggest that antibiotic-induced sliding motility results from crosstalk of engaged metabolic pathways governed by a network of regulators. While the mechanical requirements (such as surfactin, EPS, K^+^, BslA, TasA, and BCFAs) are likely the same for *B. subtilis* sliding motility triggered by different environmental cues, the specific regulatory mechanisms vary based on the initiating conditions (i.e. semi-solid media, nutrient availability, antibiotic-suppressed translation). Comparable to multicellular organisms where differentiated cells perform specific functions, heterogeneity in bacterial populations enables cells in different metabolic states to communicate and exchange information (Chou et al., 2022; Liu et al., 2015; Pisithkul et al., 2019). The detailed mechanism that initiates colony mobilization awaits further investigation to identify the specific functions connecting translation and regulatory network activation for sliding motility. Additional inducer metabolites and possibly new mechanisms of induction may reveal many possible signals that converge on a conserved response to mobilize a population under competition.

## Methods

### Strains, primers and growth media

The strains of *Bacillus subtilis* used in this study are listed in **Table S4**. *Bacillus subtilis* mutant strains in 168 or PY79 background were transduced to NCIB3610 by SPP1 phage transduction (Yasbin and Young, 1974). The primers are listed in **Table S5**. *Bacillus subtilis* strains were cultured at 37℃ in lysogeny broth (LB) and were inoculated onto GYM7 plates (0.4% [w/v] D-glucose, 0.4% [w/v] yeast extract, 1.0% [w/v] malt extract, pH7.0) with 1.5% [w/v] agar, MSgg plates (5 mM potassium phosphate buffer, diluted from 0.5 M stock with 2.72 g K_2_HPO_4_ and 1.275 g KH_2_PO_4_, pH 7.0 in 50 mL; 100 mM MOPS buffer, pH 7.0, adjusted with NaOH; 2 mM MgCl_2_•6H_2_O; 700 mM CaCl_2_•2H_2_O; 100 mM FeCl_3_•6H_2_O; 50 mM MnCl_2_•4H_2_O; 1 mM ZnCl_2_; 2 mM thiamine HCl; 0.5% [v/v] glycerol; and 0.5% [w/v] monosodium glutamate) and CM plates (Fall et al., 2004) when grown to an OD600 of 1.0. *Streptomyces venezuelae* spore stock was maintained in sterile water at 4℃.

### Construction of P*pdhA*-*lux* and P*pucA*-*lux* reporter strains

To construct luciferase reporter strains, we used primers for each promoter region (Pair 1: P*pdhA*-For and P*pdhA*-Rev; Pair 2: P*pucA*-For and P*pucA*-Rev) to amplify the target promoter from *B. subtilis* NCIB 3610 genomic DNA, primers *lux*-For and *lux*-Rev to amplify the *luxABCDE* fragment from pBS3Klux, primers *amyE*-back-For and *amyE*-front-Rev to amplify the plasmid backbone (including the origin site, ampicillin resistance cassette, and front and back partial *amyE* fragments) from pDR111, and primers *kan*-For and *kan*-Rev to amplify the kanamycin resistance cassette from pDG780 (Stubbendieck and Straight, 2015). These 4 fragments will be assembled to a functional plasmid by Gibson assembly (Gibson et al., 2009). The plasmid was transformed to *B. subtilis* PY79 wild type and then the construct containing the target promoter, *lux* operon, and kanamycin resistance cassette was inserted into the *amyE* locus. The inserted construct was verified by PCR. Once confirmed, the construct was moved to PDS0066 (*B. subtilis* NCIB 3610 wild type) and PDS0972 (*B. subtilis* NCIB 3610 Δ*codY*) using SPP1 phage transduction to generate PDS0981, PDS0982, PDS0983, and PDS0984.

### Sliding motility assay

*B. subtilis* cells grown in 5 ml LB broth were diluted to an OD600 of 0.08. When grown to an OD600 of 1.0, 1.5 μL of *B. subtilis* cells were spotted on the GYM7 plate with or without 1 μM chloramphenicol. Pictures were taken with Nikon D60 digital camera. For the luciferase reporter assays, images were captured with an Amersham imager 600, or Canon 5D Mark IV (Brake et al., 2021). For the coculture assay, 2.5 μL of *Streptomyces venezuelae* spores (10^7^ spores/mL) and 1.5 μL of *B. subtilis* (OD600=1.0, equal to 10^8^ cells/mL) cells were spotted in a cross pattern with a distance of 1 cm between spots for each species.

### RNA extraction

Wild-type and Δ*codY B. subtilis* NCIB3610 were grown to early stationary phase (OD600=1.0) and inoculated on GYM7 plates without or with 1 μM chloramphenicol, followed by incubation at 30 ℃ for 6h and 24h. *Bacillus subtilis* colonies at 6 h and 24 h were scraped from agar plates. RNA was extracted using TRI reagent (Sigma) with standard procedures (Stubbendieck and Straight, 2017). Turbo DNA-free kit (Applied Biosystems) was used to remove DNA from RNA samples according to the standard protocol.

### RNA-Seq and analysis

50 bp single-end read libraries were constructed using a TruSeq Stranded mRNA Kit and sequenced using the Illumina HiSeq2500. We generated raw counts by mapping reads to all the open reading frames (ORFs) in the *B. subtilis* 168 genome (Gene Bank: NC_000964.3) plus *sfp*, *swrA* and ORFs in the plasmid pSB32 with Kallisto (V0.48.0) (Bray et al., 2016). Raw counts were used as the input for DESeq2 (R/Bioconductor) (Love et al., 2014) to generate normalized counts for differentially gene expression analysis. Transcript data were normalized to library size using RNAlysis (Teichman et al., 2023)(V3.7.0). ORFs with |fold change|>=2 and Padj<0.05 are defined as differentially expressed genes in this study. The raw reads in this study are accessible from NCBI BioProject Accession GSE261934.

### Cytoscape network

Identities of known regulons were downloaded from Subtiwiki (Zhu and Stülke, 2018) and paired with our transcriptomic data. The output was organized as regulatory networks that emerge at 6 and 24 hours based on the number of genes within a given regulon that change expression by ≥ 2X and adjusted p<0.05. Each node represents an individual regulator. Node size represents the normalized number of genes that change expression relative to the number of overall genes in each individual regulon. The edge connecting nodes represents shared genes between two regulons. Edge width represents the normalized number of shared genes that change expression relative to the number of shared genes between two regulons. Blue and red inside each node represent the percentage of upregulation and downregulation in each regulon, respectively.

### Metabolite extraction

Metabolite extraction was adapted from the previous method (Rosebrock and Caudy, 2017). Briefly, cells were collected with cell scrapers from agar plates and immediately transferred to low-temperature organic solvent (methanol: acetonitrile: H2O=40:40:20). After three rounds of freeze-thaws between −20 °C and −80 °C, cells were spun down and saved for DNA content measurement by diphenylamine assay and supernatants were collected.

Samples were dried under N2 gas and reconstituted in high-purity water according to optical density at harvest. Reconstituted samples were combined with an equal volume of 13C/15N internal standard mixture composed of 99.9% 13C15N-labeled *Saccharomyces cerevisiae* extract isolated from a range of culture conditions. Following mixture with internal standard, samples were cleared for 7 min at 14,000 × *g* at 4 °C and transferred to LC-MS polypropylene conical vials (Agilent). Vials were stored at −80 °C until analysis using LC-MS.

#### Chromatographic separation

All metabolomics samples described in this study were analyzed by chromatographic methods essentially as described in Caudy et al. (Caudy et al., 2019). Briefly, tributylamine ion paired LC was carried out using an Agilent 1290 UPLC system with a 0.25 mL/min flow rate and a 2-μL sample loop and an Extend C18 RRHD 1.8 μm, 2.1 × 150 mm column (Agilent).

#### Mass spectrometry

Data were acquired using an Agilent 6550 QToF instrument. Samples were ionized using an Agilent Jet Stream electrospray ionization source operated in negative mode. Gas temperature in the ion source was 150 °C with a flow rate of 14 L/min. Nebulizer pressure was 45 psig; sheath temperature was 325 °C with a gas flow rate of 12 L/min. Voltage for both capillary and nozzle was 2000 V. The funnel DC voltage was −30 V, funnel voltage drops were −100 and −50 V in the high- and low-pressure funnels, respectively. RF voltages were 110 and 60 V in the high- and low-pressure funnels respectively. Mass spectra were acquired between 50 *m*/*z* and 1100 *m*/*z* with a rate of 2 spectra per second. Mass lock mixture was used as described in (Caudy et al., 2019).

#### Data analysis

Quantitation of each analyte was carried out by calculating the area under the curve of a chromatographic trace corresponding to the [M-H]− ion of the analyte of interest and its 13C/15N-fully labeled variant. Ion chromatograms were extracted from raw profile mass spectra using chromXtractorPro, an R package developed in the Rosebrock laboratory (available on request). This software was used to set consistent margins for peak integration by locally aligning and manually integrating chromatograms corresponding to extracted mass windows of 75 ppm around each reference mass. Retention times for each analyte were determined using pure standards.

### Click-iT assay for protein synthesis

The method used in this study was adapted from Click-iT HPG Alexa Fluor protein synthesis assay with slight modifications. *B. subtilis* cells cultured in S7 (methionine-free) medium overnight were passaged to 500 μL S7 medium to an optical density of 0.08 at 600 nm (OD600=0.08) and grown to OD600=0.5 in a 30 °C shaker with a speed of 230 rpm in the dark. 5 μL of 100 μM Cm was added to 495 μL *B. subtilis* culture at OD600=0.08 to a final concentration of 1 μM Cm. Then, 5 μL of 5 mM HPG was added to the culture to a final concentration of 50 μM and grown in the same condition. For, 16 μM Cm sample, 5 μL of 1.6 mM Cm was added to 490 μL of *B. subtilis* culture with 5 μL of 5 mM HPG at OD600=0.5 to a final concentration of 16 μM Cm. After 5 min incubation, S7 medium containing HPG was removed (Eppendorf benchtop centrifuge, 15,000 rpm, 1 minute) and cells were washed once with 500 μL of PBS (15,000 rpm, 1 minute). Then, cells were fixed in 500 μL of 4% paraformaldehyde (PFA) (in 1X PBS) at room temperature for 7 minutes. Cells were washed once with 500 μL of PBS and resuspended in 500 μL of PBS followed by sonication for 20 s to separate cells (1 s/1 s; 20% amplitude with Thermo FB-20). For permeabilization, cells were washed twice with 3% BSA (in 1X PBS), and then permeabilized in 500 μL of 0.5% Triton X-100 (in 1X PBS) for 20 minutes at room temperature. To detect the HPG incorporation, we followed the Click-iT HPG Alexa Fluor protein synthesis assay kits user guide. Briefly, cells were washed twice with 3% BSA (in 1X PBS) to remove the permeabilization buffer and resuspended in 500 μL of Click-iT reaction cocktail (prepared 15 minutes before the reaction) for a total of 30-minute incubation in the dark. Cells were further washed with Click-iT reaction rinse buffer and resuspended in PBS before flow cytometry analysis (BD Accuri C6). The mean fluorescence intensity was calculated with 20,000 cells for each sample.

## Acknowledgements

We thank Reed Stubbendieck, Sandra Truong, and Ryan McCormick for the discussion on the RNA-Seq data analysis and Kailu Yang for the discussion on the Cytoscape network analysis. We thank Abraham Sonenshein and Lianet Noda-Garcia for helpful comments on the manuscript. Research reported in this publication was supported by the National Institute of General Medical Sciences of the National Institutes of Health under Award Number R01GM141700 and in part by an NSF-CAREER Award (MCB-1253215). The content is solely the responsibility of the authors and does not necessarily represent the official views of the National Institutes of Health or National Science Foundation.

---

**Figure S1.**
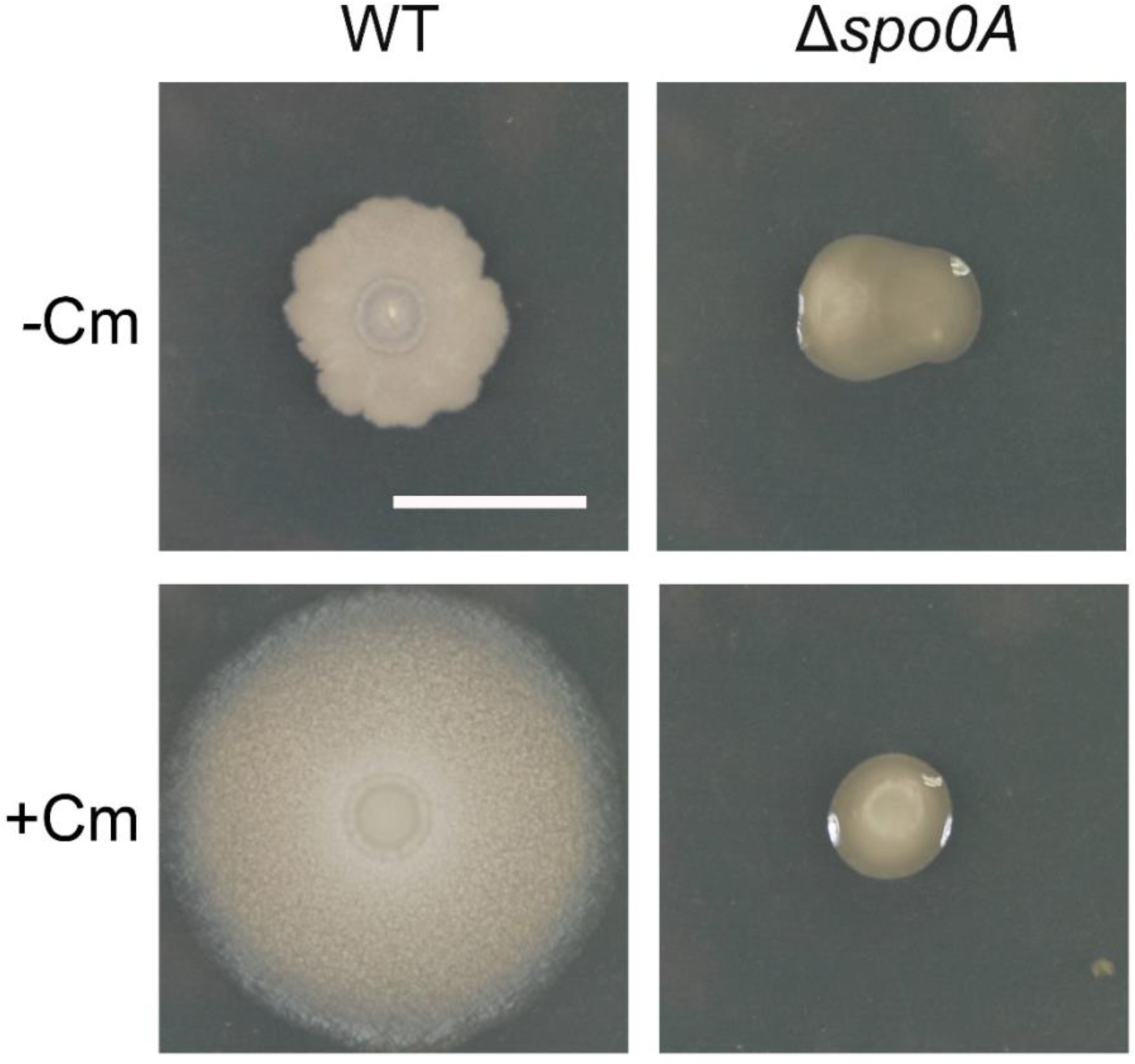
The colony of Δ*spo0A* strain was unable to expand in the presence of chloramphenicol. Wild type (WT) and Δ*spo0A B. subtilis* NCIB3610 were spotted on the GYM7 plate in the absence (-Cm) and presence (+Cm) of chloramphenicol. Pictures were taken at 24 h. Bar, 1 cm.

**Figure S2.**
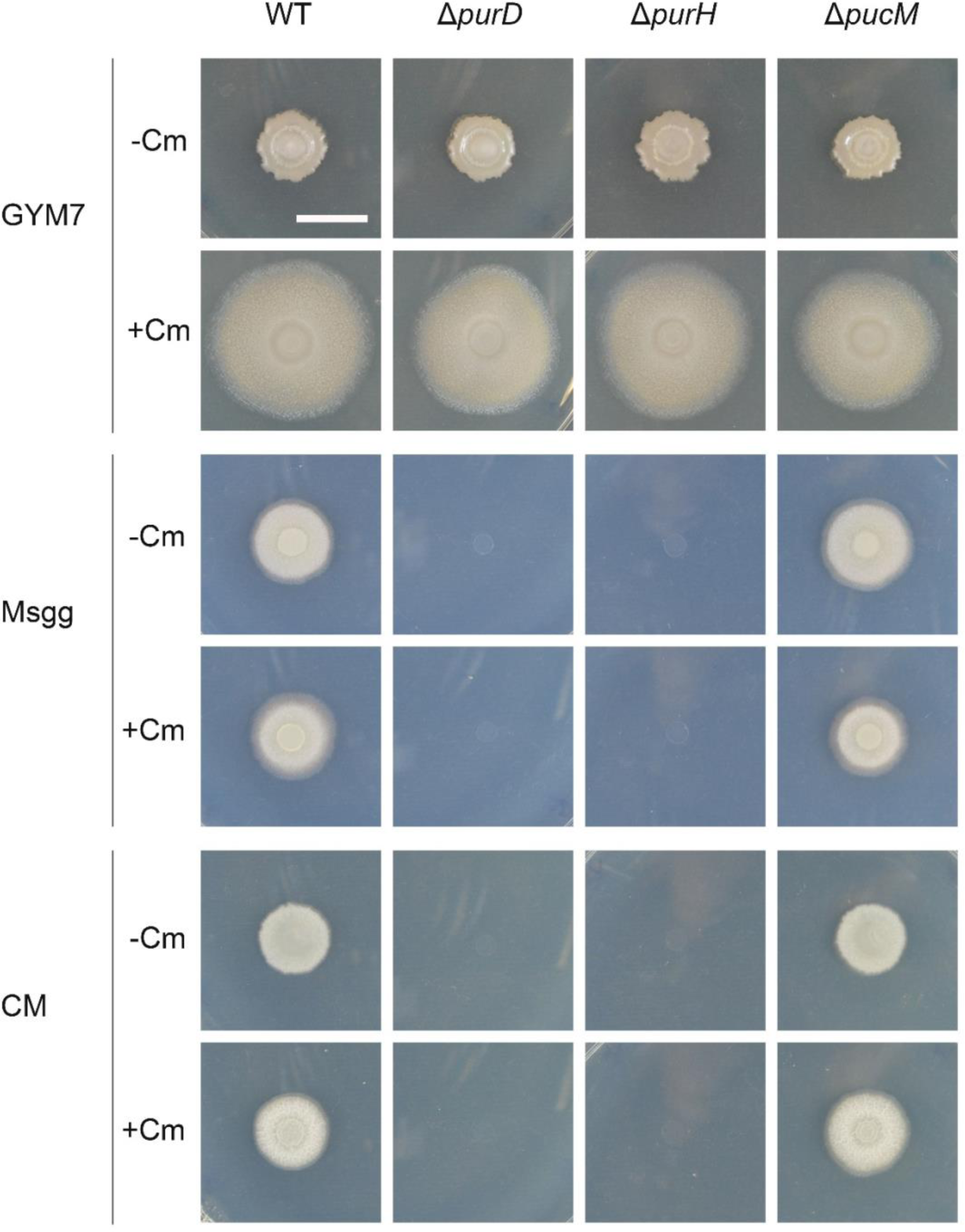
Disruption of purine de novo biosynthesis pathway or catabolism pathway has different effects on *B. subtilis* growth under different conditions. Wild type, Δ*purD*, Δ*purH*, and Δ*purM B. subtilis* NCIB3610 were spotted on the agar plate (1.5% agar) with GYM7 (top), Msgg (middle), and CM (bottom) media in the absence (-Cm) and presence (+Cm) of chloramphenicol. Pictures were taken at 24 h. Bar, 1 cm.

**Figure S3.**
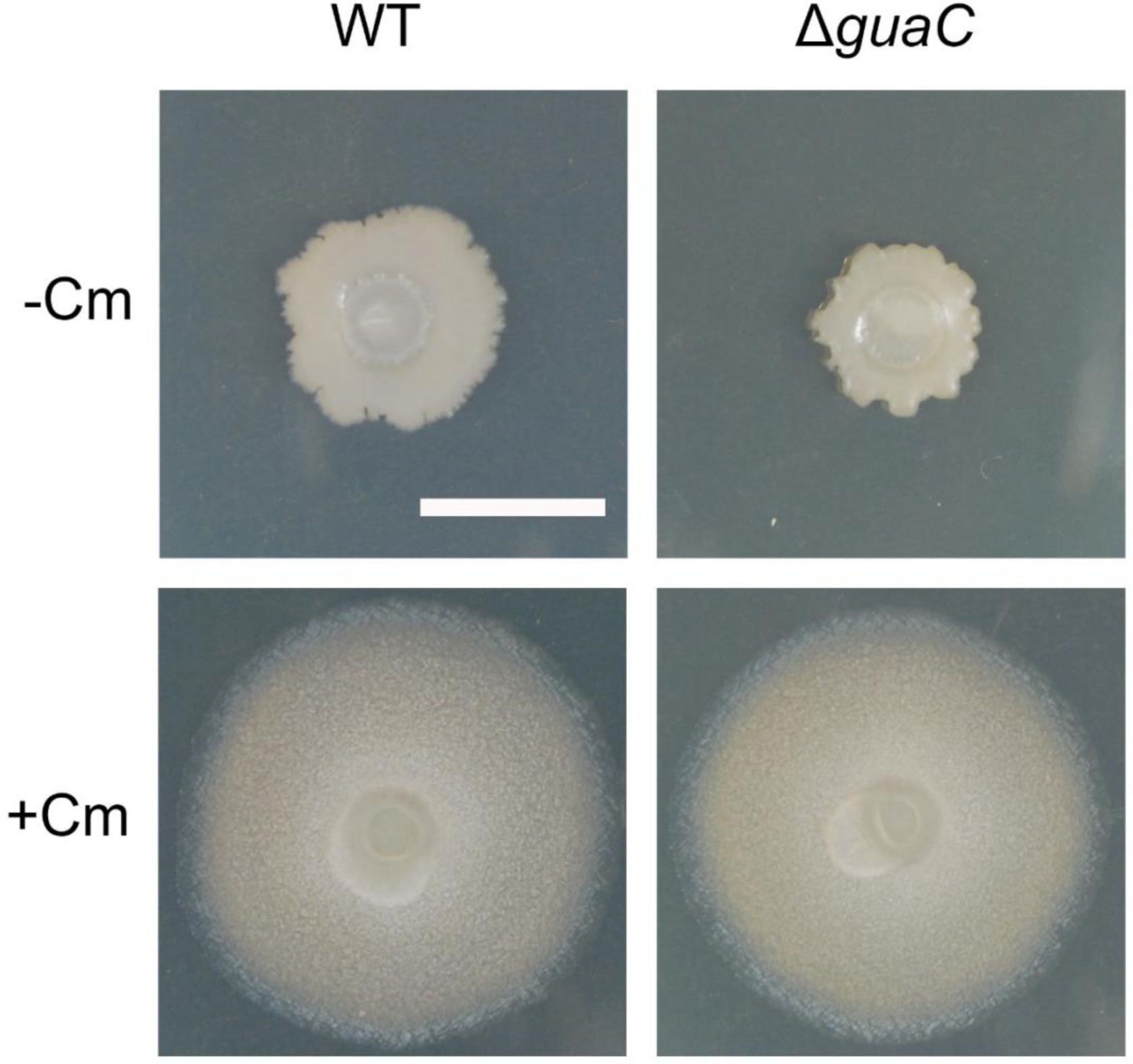
The colony of Δ*guaC* strain was able to expand in the presence of chloramphenicol. Wild type (WT) and Δ*guaC B. subtilis* NCIB3610 were spotted on the GYM7 plate in the absence (-Cm) and presence (+Cm) of chloramphenicol. Pictures were taken at 24 h. Bar, 1 cm.

**Figure S4.**
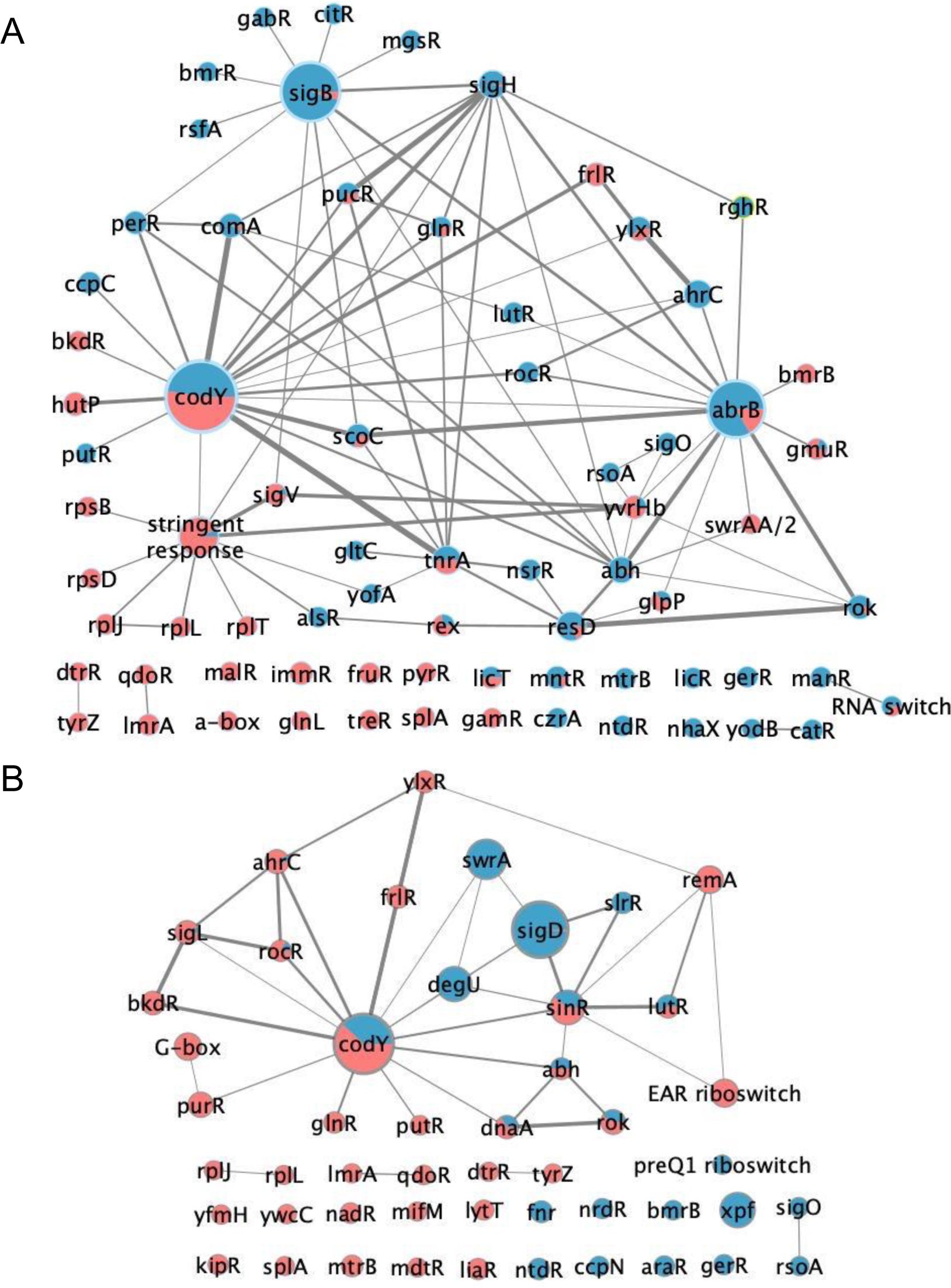
A network view *ΔcodY* regulation in a constitutively mobile population. Cytoscape networks of regulators with ≥ 40% engagement of regulated genes at either: **A.** 6 h *ΔcodY* gene expression ≥ 2-fold difference from wildtype controls with adjusted p-value < 0.05, or **B.** 24 h *ΔcodY* gene expression ≥ 2-fold difference from untreated controls with adjusted p-value < 0.05. In both A and B, the size of each node represents weighted gene number in each regulon (weighted by percentage of differentially expressed genes in each regulon) and the edge width represents weighted overlapped gene number between two regulons (weighted by percentage of differentially expressed genes among overlapped genes). Blue: downregulation; Red: upregulation.

**Figure S5.**
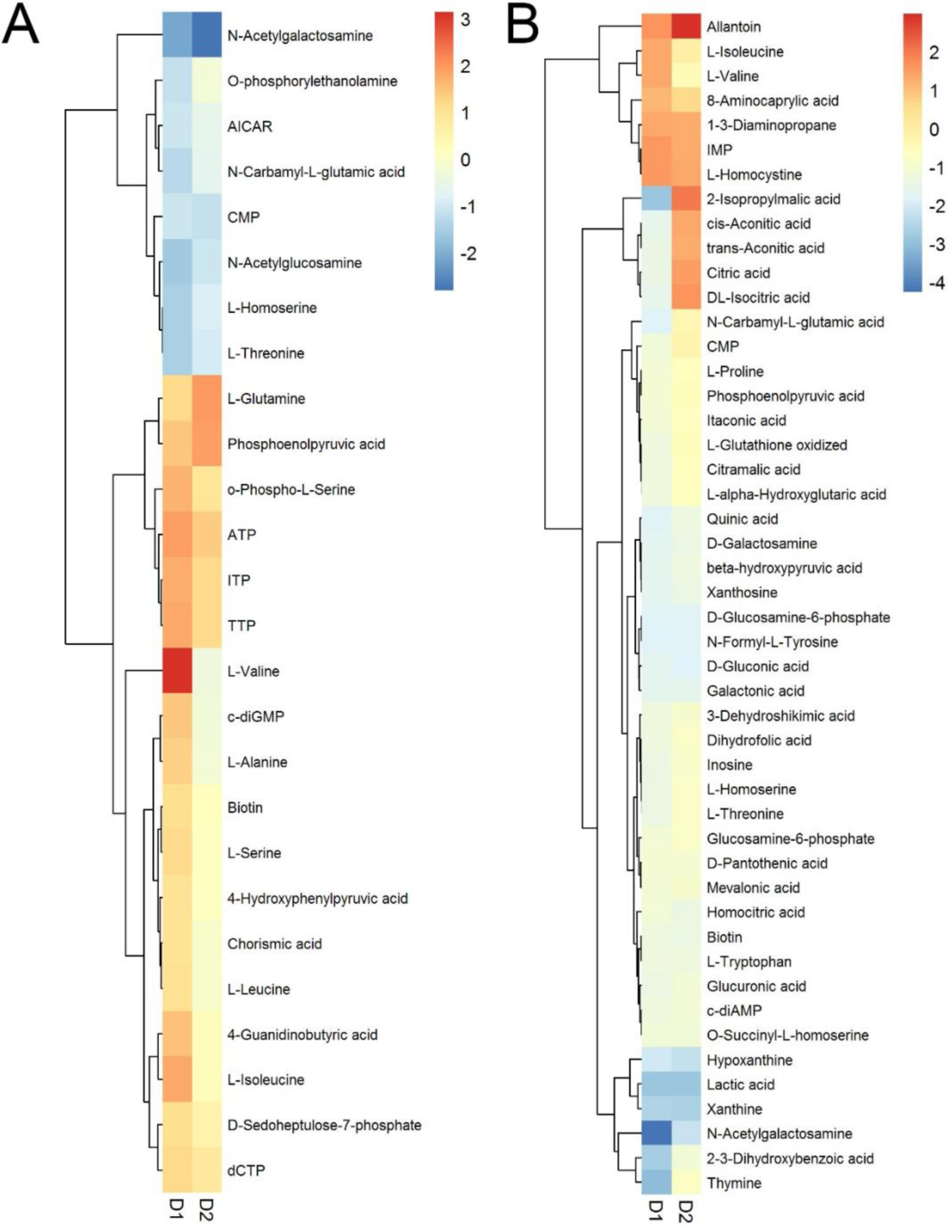
Metabolomics analysis underscores the pattern of shifting metabolism reflected by transcriptional analysis. The same data used for Figure 7 is presented here with Δ*codY* as the D1 and Cm-treated as the D2 for comparison. Metabolic profiles of Δ*codY* (D1) strain at 6 h (**A**) and 24 h (**B**). Metabolites that change ≥1.5-fold (6 h) and ≥2-fold (24 h) in Δ*codY* compared with wild-type strain are listed. The profiles of corresponding metabolites in wild-type strain (D2) upon exposure to 1 μM Cm at 6 h and 24 h.

**Figure S6.**
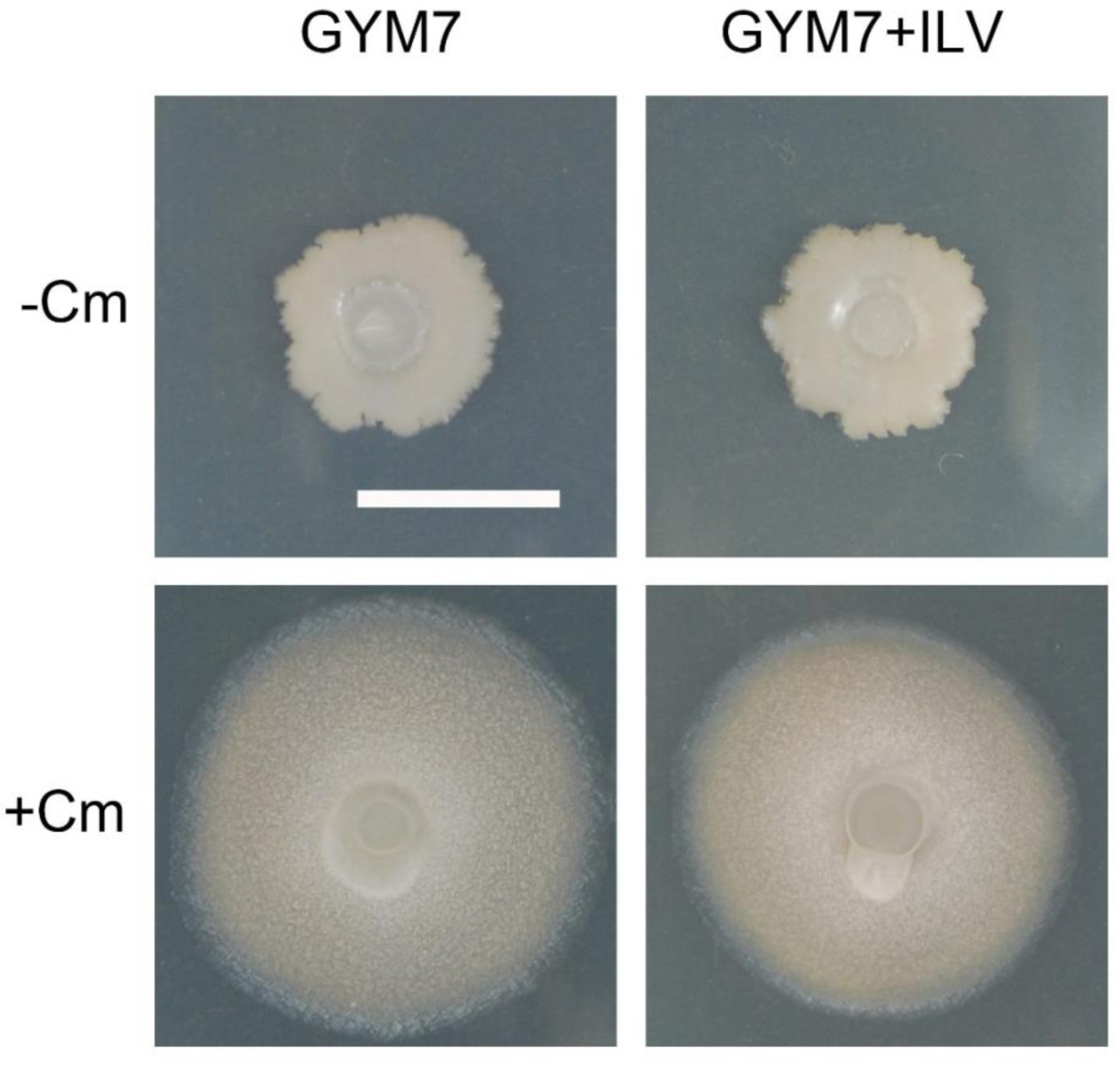
Supplementation of BCAAs was unable to inhibit sliding motility in the presence of chloramphenicol. Wild type B. subtilis NCIB3610 was spotted on the GYM7 plate and GYM7 supplemented with 10 mM each of isoleucine, leucine and valine in the absence (-Cm) and presence (+Cm) of chloramphenicol. Pictures were taken at 24 h. Bar, 1 cm.

## References

Alves, M., Al-badi, E., Withycombe, C., Jones, P.M., Purdy, K.J., Maddocks, S.E., 2018. Interaction between Staphylococcus aureus and Pseudomonas aeruginosa is beneficial for colonisation and pathogenicity in a mixed biofilm. FEMS 76, 1–10. 10.1093/femspd/fty003

Bader, M.W., Navarre, W.W., Shiau, W., Nikaido, H., Frye, J.G., McClelland, M., Fang, F.C., Miller, S.I., 2003. Regulation of Salmonella typhimurium virulence gene expression by cationic antimicrobial peptides. Mol. Microbiol. 50, 219–230. 10.1046/j.1365-2958.2003.03675.x

Banse, A.V., Chastanet, A., Rahn-Lee, L., Hobbs, E.C., Losick, R., 2008. Parallel pathways of repression and antirepression governing the transition to stationary phase in Bacillus subtilis. Proc. Natl. Acad. Sci. 105, 15547–15552. 10.1073/pnas.0805203105

Basler, M., Ho, B.T., Mekalanos, J.J., 2013. Tit-for-Tat: Type VI Secretion System Counterattack during Bacterial Cell-Cell Interactions. Cell 152, 884–894. 10.1016/j.cell.2013.01.042

Beier, L., Nygaard, P., Jarmer, H., Saxild, H.H., 2002. Transcription analysis of the Bacillus subtilis PucR regulon and identification of a cis-acting sequence required for PucR-regulated expression of genes involved in purine catabolism. J. Bacteriol. 184, 3232– 3241. 10.1128/JB.184.12.3232-3241.2002

Binnie, C., Lampe, M., Losick, R., 1986. Gene encoding the sigma 37 species of RNA polymerase sigma factor from Bacillus subtilis. Proc. Natl. Acad. Sci. U. S. A. 83, 5943– 5947. 10.1073/pnas.83.16.5943

Bosdriesz, E., Molenaar, D., Teusink, B., Bruggeman, F.J., 2015. How fast-growing bacteria robustly tune their ribosome concentration to approximate growth-rate maximization. FEBS J. 282, 2029–2044. 10.1111/febs.13258

Brake, C., Liu, Y., Straight, P., LaFayette, C., 2021. MicroWorlds: A macro study of microbial interactions informs a bio-art series, in: 2021 25th International Conference Information Visualisation (IV). Presented at the 2021 25th International Conference Information Visualisation (IV), pp. 297–302. 10.1109/IV53921.2021.00055

Bray, N.L., Pimentel, H., Melsted, P., Pachter, L., 2016. Near-optimal probabilistic RNA-seq quantification. Nat. Biotechnol. 34, 525–527. 10.1038/nbt.3519

Brien, J.O., Wright, G.D., 2011. An ecological perspective of microbial secondary metabolism. Curr. Opin. Biotechnol. 22, 552–558. 10.1016/j.copbio.2011.03.010

Brinsmade, S.R., 2016. CodY, a master integrator of metabolism and virulence in Gram-positive bacteria. Curr. Genet. 1–9. 10.1007/s00294-016-0656-5

Brinsmade, S.R., Alexander, E.L., Livny, J., Stettner, A.I., Segrè, D., Rhee, K.Y., Sonenshein, A.L., 2014. Hierarchical expression of genes controlled by the Bacillus subtilis global regulatory protein CodY. Proc. Natl. Acad. Sci. U. S. A. 111, 2–7. 10.1073/pnas.1321308111

Burbulys, D., Trach, K.A., Hoch, J.A., 1991. Initiation of sporulation in B. subtilis is controlled by a multicomponent phosphorelay. Cell 64, 545–552. 10.1016/0092-8674(91)90238-T

Cao, H., Kuipers, O.P., 2018. Influence of global gene regulatory networks on single cell heterogeneity of green fluorescent protein production in Bacillus subtilis. Microb. Cell Factories 17, 134. 10.1186/s12934-018-0985-9

Caudy, A.A., Hanchard, J.A., Hsieh, A., Shaan, S., Rosebrock, A.P., 2019. Functional genetic discovery of enzymes using full-scan mass spectrometry metabolomics (1). Biochem. Cell Biol. Biochim. Biol. Cell. 97, 73–84. 10.1139/bcb-2018-0058

Chou, K.-T., Lee, D.D., Chiou, J., Galera-Laporta, L., Ly, S., Garcia-Ojalvo, J., Süel, G.M., 2022. A segmentation clock patterns cellular differentiation in a bacterial biofilm. Cell 185, 145–157.e13. 10.1016/j.cell.2021.12.001

Chumsakul, O., Nakamura, K., Kurata, T., Sakamoto, T., Hobman, J.L., Ogasawara, N., Oshima, T., Ishikawa, S., 2013. High-Resolution Mapping of In vivo Genomic Transcription Factor Binding Sites Using In situ DNase I Footprinting and ChIP-seq. DNA Res. 20, 325–338. 10.1093/dnares/dst013

Dar, D., Shamir, M., Mellin, J.R., Koutero, M., Stern-Ginossar, N., Cossart, P., Sorek, R., 2016. Term-seq reveals abundant ribo-regulation of antibiotics resistance in bacteria. Science 352, aad9822. 10.1126/science.aad9822

Davies, J., 2006. Are antibiotics naturally antibiotics ? J Ind Microbiol Biotechnol 33, 496–499. 10.1007/s10295-006-0112-5

Davies, J., Spiegelman, G.B., Yim, G., 2006. The world of subinhibitory antibiotic concentrations. Curr. Opin. Microbiol. 9, 445–453. 10.1016/j.mib.2006.08.006

Dubnau, D., Losick, R., 2006. Bistability in bacteria. Mol. Microbiol. 61, 564–572.

Eldar, A., Chary, V.K., Xenopoulos, P., Fontes, M.E., Losón, O.C., Dworkin, J., Piggot, P.J., Elowitz, M.B., 2009. Partial penetrance facilitates developmental evolution in bacteria. Nature 460, 510–514. 10.1038/nature08150

Endo, T., Uratani, B., Freese, E., 1983. Purine salvage pathways of Bacillus subtilis and effect of guanine on growth of GMP reductase mutants. J. Bacteriol. 155, 169–179. 10.1128/jb.155.1.169-179.1983

Erez, A., Kolodkin-Gal, I., 2017. From Prokaryotes to Cancer: Glutamine Flux in Multicellular Units. Trends Endocrinol. Metab. 28, 637–644. 10.1016/j.tem.2017.05.007

Fajardo, A., Martínez, J.L., 2008. Antibiotics as signals that trigger specific bacterial responses. Curr. Opin. Microbiol. 11, 161–167. 10.1016/j.mib.2008.02.006

Fall, R., Kinsinger, R.F., Wheeler, K.A., 2004. A Simple Method to Isolate Biofilm-forming Bacillus subtilis and Related Species from Plant Roots. Syst. Appl. Microbiol. 27, 372–9.

Fujita, M., González-Pastor, J.E., Losick, R., 2005. High- and low-threshold genes in the Spo0A regulon of Bacillus subtilis. J. Bacteriol. 187, 1357–1368. 10.1128/JB.187.4.1357-1368.2005

Galet, J., Deveau, A., Hôtel, L., Frey-Klett, P., Leblond, P., Aigle, B., 2015. Pseudomonas fluorescens pirates both ferrioxamine and ferricoelichelin Siderophores from Streptomyces ambofaciens. Appl. Environ. Microbiol. 81, 3132–3141. 10.1128/AEM.03520-14

Gibson, D.G., Young, L., Chuang, R.-Y., Venter, J.C., Hutchison, C.A., Smith, H.O., 2009. Enzymatic assembly of DNA molecules up to several hundred kilobases. Nat. Methods 6, 343–345. 10.1038/nmeth.1318

Goh, E.B., Yim, G., Tsui, W., McClure, J., Surette, M.G., Davies, J., 2002. Transcriptional modulation of bacterial gene expression by subinhibitory concentrations of antibiotics. Proc. Natl. Acad. Sci. U. S. A. 99, 17025–17030. 10.1073/pnas.252607699

Grau, R.R., de Oña, P., Kunert, M., Leñini, C., Gallegos-Monterrosa, R., Mhatre, E., Vileta, D., Donato, V., Hölscher, T., Boland, W., Kuipers, O.P., Kovács, Á.T., 2015. A Duo of Potassium-Responsive Histidine Kinases Govern the Multicellular Destiny of Bacillus subtilis. mBio 6, e00581. 10.1128/mBio.00581-15

Groisman, E.A., Chan, C., 2021. Cellular Adaptations to Cytoplasmic Mg2+ Limitation. Annu. Rev. Microbiol. 75, 649–672. 10.1146/annurev-micro-020518-115606

Grundy, F.J., Waters, D.A., Allen, S.H., Henkin, T.M., 1993. Regulation of the Bacillus subtilis acetate kinase gene by CcpA. J. Bacteriol. 175, 7348–7355. 10.1128/jb.175.22.7348-7355.1993

Gyan, S., Shiohira, Y., Sato, I., Takeuchi, M., Sato, T., 2006. Regulatory Loop between Redox Sensing of the NADH/NAD+ Ratio by Rex (YdiH) and Oxidation of NADH by NADH Dehydrogenase Ndh in Bacillus subtilis. J. Bacteriol. 188, 7062–7071. 10.1128/jb.00601-06

Haldenwang, W.G., Losick, R., 1979. A modified RNA polymerase transcribes a cloned gene under sporulation control in Bacillus subtilis. Nature 282, 256–260. 10.1038/282256a0

Harrison, F., Paul, J., Massey, R.C., Buckling, A., 2008. Interspecific competition and siderophore-mediated cooperation in Pseudomonas aeruginosa. ISME J. 2, 49–55. 10.1038/ismej.2007.96

Hauryliuk, V., Atkinson, G.C., Murakami, K.S., Tenson, T., Gerdes, K., 2015. Recent functional insights into the role of (p)ppGpp in bacterial physiology. Nat. Rev. Microbiol. 13, 298–309. 10.1038/nrmicro3448

Hecker, M., Pané-Farré, J., Völker, U., 2007. SigB-dependent general stress response in Bacillus subtilis and related gram-positive bacteria. Annu. Rev. Microbiol. 61, 215–236. 10.1146/annurev.micro.61.080706.093445

Hecker, M., Völker, U., 1998. Non-specific, general and multiple stress resistance of growth-restricted Bacillus subtilis cells by the expression of the σ(B) regulon. Mol. Microbiol. 29, 1129–1136. 10.1046/j.1365-2958.1998.00977.x

Hernandez-Valdes, J.A., Zhou, L., de Vries, M.P., Kuipers, O.P., 2020. Impact of spatial proximity on territoriality among human skin bacteria. Npj Biofilms Microbiomes 6, 1–13. 10.1038/s41522-020-00140-0

Herold, S., Siebert, J., Huber, A., Schmidt, H., 2005. Global expression of prophage genes in Escherichia coli O157:H7 strain EDL933 in response to norfloxacin. Antimicrob. Agents Chemother. 49, 931–944. 10.1128/AAC.49.3.931-944.2005

Hoefler, B.C., Gorzelnik, K. V., Yang, J.Y., Hendricks, N., Dorrestein, P.C., Straight, P.D., 2012. Enzymatic resistance to the lipopeptide surfactin as identified through imaging mass spectrometry of bacterial competition. Proc. Natl. Acad. Sci. 109, 13082–13087. 10.1073/pnas.1205586109

Hogan, D.A., Kolter, R., 2002. Pseudomonas-Candida Interactions: An Ecological Role for Virulence Factors. Science 296, 2229–2232. 10.1126/science.1070784

Hölscher, T., Kovács, Á.T., 2017. Sliding on the surface: bacterial spreading without an active motor. Environ. Microbiol. 19, 2537–2545. 10.1111/1462-2920.13741

Jacobson, A., Gillespie, D., 1968. Metabolic events occurring during recovery from prolonged glucose starvation in Escherichia coli. J. Bacteriol. 95, 1030–1039. 10.1128/jb.95.3.1030-1039.1968

Jeckel, H., Nosho, K., Neuhaus, K., Hastewell, A.D., Skinner, D.J., Saha, D., Netter, N., Paczia, N., Dunkel, J., Drescher, K., 2023. Simultaneous spatiotemporal transcriptomics and microscopy of Bacillus subtilis swarm development reveal cooperation across generations. Nat. Microbiol. 8, 2378–2391. 10.1038/s41564-023-01518-4

Jones, S.E., Ho, L., Rees, C.A., Hill, J.E., Nodwell, J.R., Elliot, M.A., 2017. Streptomyces exploration is triggered by fungal interactions and volatile signals. eLife 6, e21738. 10.7554/eLife.21738

Julkowska, D., Obuchowski, M., Holland, I.B., Séror, S.J., 2005. Comparative analysis of the development of swarming communities of Bacillus subtilis 168 and a natural wild type: critical effects of surfactin and the composition of the medium. J. Bacteriol. 187, 65–76. 10.1128/JB.187.1.65-76.2005

Kearns, D.B., 2010. A field guide to bacterial swarming motility. Nat. Rev. Microbiol. 8, 634–644. 10.1038/nrmicro2405

Kearns, D.B., Chu, F., Branda, S.S., Kolter, R., Losick, R., 2005. A master regulator for biofilm formation by Bacillus subtilis. Mol. Microbiol. 55, 739–749. 10.1111/j.1365-2958.2004.04440.x

Kearns, D.B., Losick, R., 2004. Swarming motility in undomesticated Bacillus subtilis. Mol. Microbiol. 49, 581–590. 10.1046/j.1365-2958.2003.03584.x

Kinsinger, R.F., Kearns, D.B., Hale, M., Fall, R., 2005. Genetic Requirements for Potassium Ion-Dependent Colony Spreading in Bacillus subtilis. J. Bacteriol. 187, 8462–8469. 10.1128/JB.187.24.8462-8469.2005

Kinsinger, R.F., Shirk, M.C., Fall, R., 2003. Rapid Surface Motility in Bacillus subtilis Is Dependent on Extracellular Surfactin and Potassium Ion. J. Bacteriol. 185, 5627–5631. 10.1128/JB.185.18.5627-5631.2003

Kovács, Á.T., 2016. Bacterial differentiation via gradual activation of global regulators. Curr. Genet. 62, 125–8. 10.1007/s00294-015-0524-8

Kriel, A., Bittner, A.N., Kim, S.H., Liu, K., Tehranchi, A.K., Zou, W.Y., Rendon, S., Chen, R., Tu, B.P., Wang, J.D., 2012. Direct regulation of GTP homeostasis by (p)ppGpp: A critical component of viability and stress resistance. Mol. Cell 48, 231–241. 10.1016/j.molcel.2012.08.009

Kriel, A., Brinsmade, S.R., Tse, J.L., Tehranchi, A.K., Bittner, A.N., Sonenshein, A.L., Wang, J.D., 2014. GTP Dysregulation in Bacillus subtilis Cells Lacking (p)ppGpp Results in Phenotypic Amino Acid Auxotrophy and Failure To Adapt to Nutrient Downshift and Regulate Biosynthesis Genes. J. Bacteriol. 196, 189–201. 10.1128/JB.00918-13

Kukurugya, M.A., Titov, D.V., 2023. The Warburg Effect is the result of faster ATP production by glycolysis than respiration. 10.1101/2022.12.28.522160

LeRoux, M., Kirkpatrick, R.L., Montauti, E.I., Tran, B.Q., Peterson, S.B., Harding, B.N., Whitney, J.C., Russell, A.B., Traxler, B., Goo, Y.A., Goodlett, D.R., Wiggins, P.A., Mougous, J.D., 2015. Kin cell lysis is a danger signal that activates antibacterial pathways of Pseudomonas aeruginosa. eLife 4, e05701. 10.7554/eLife.05701

Lin, J.T., Connelly, M.B., Amolo, C., Otani, S., Yaver, D.S., 2005. Global Transcriptional Response of Bacillus subtilis to Treatment with Subinhibitory Concentrations of Antibiotics That Inhibit Protein Synthesis. Antimicrob. Agents Chemother. 49, 1915– 1926. 10.1128/AAC.49.5.1915-1926.2005

Linares, J.F., Gustafsson, I., Baquero, F., Martinez, J.L., 2006. Antibiotics as intermicrobiol signaling agents instead of weapons. Proc. Natl. Acad. Sci. U. S. A. 103, 19484–19489. 10.1073/pnas.0608949103

Liu, J., Prindle, A., Humphries, J., Gabalda-Sagarra, M., Asally, M., Lee, D.Y.D., Ly, S., Garcia-Ojalvo, J., Süel, G.M., 2015. Metabolic co-dependence gives rise to collective oscillations within biofilms. Nature 523, 550–554. 10.1038/nature14660

Liu, Y., Kyle, S., Straight, P.D., 2018. Antibiotic Stimulation of a Bacillus subtilis Migratory Response. mSphere 3, e00586–17. 10.1128/mSphere.00586-17

López, D., Kolter, R., 2010. Extracellular signals that define distinct and coexisting cell fates in Bacillus subtilis. FEMS Microbiol. Rev. 34, 134–149. 10.1111/j.1574-6976.2009.00199.x

Lopez, D., Vlamakis, H., Kolter, R., 2009. Generation of multiple cell types in Bacillus subtilis. FEMS Microbiol. Rev. 33, 152–163. 10.1111/j.1574-6976.2008.00148.x

Love, M.I., Huber, W., Anders, S., 2014. Moderated estimation of fold change and dispersion for RNA-seq data with DESeq2. Genome Biol. 15, 550. 10.1186/s13059-014-0550-8

Maaløe, O., 1979. Regulation of the Protein-Synthesizing Machinery—Ribosomes, tRNA, Factors, and So On, in: Goldberger, R.F. (Ed.), Biological Regulation and Development: Gene Expression, Biological Regulation and Development. Springer US, Boston, MA, pp. 487–542. 10.1007/978-1-4684-3417-0_12

Mavridou, D.A.I., Gonzalez, D., Kim, W., West, S.A., Foster, K.R., 2018. Bacteria Use Collective Behavior to Generate Diverse Combat Strategies. Curr. Biol. 28, 345–355.e4. 10.1016/j.cub.2017.12.030

McCormick, D.M., Lalanne, J.-B., Lan, T.C.T., Rouskin, S., Li, G.-W., 2021. Sigma factor dependent translational activation in Bacillus subtilis. RNA rna.078747.121. 10.1261/rna.078747.121

McCully, L.M., Bitzer, A.S., Seaton, S.C., Smith, L.M., Silby, M.W., 2019. Interspecies Social Spreading: Interaction between Two Sessile Soil Bacteria Leads to Emergence of Surface Motility. mSphere 4, 1–16. 10.1128/msphere.00696-18

McKinlay, J.B., Cook, G.M., Hards, K., 2020. Microbial energy management—A product of three broad tradeoffs, in: Advances in Microbial Physiology. Elsevier, pp. 139–185. 10.1016/bs.ampbs.2020.09.001

Mirouze, N., Prepiak, P., Dubnau, D., 2011. Fluctuations in spo0A transcription control rare developmental transitions in Bacillus subtilis. PLoS Genet. 7, e1002048. 10.1371/journal.pgen.1002048

Molle, V., Fujita, M., Jensen, S.T., Eichenberger, P., González-Pastor, J.E., Liu, J.S., Losick, R., 2003. The Spo0A regulon of Bacillus subtilis. Mol. Microbiol. 50, 1683–1701. 10.1046/j.1365-2958.2003.03818.x

Morales, D.K., Grahl, N., Okegbe, C., Dietrich, L.E.P., Jacobs, N.J., Hogan, A., 2013. Control of Candida albicans Metabolism and Biofilm Formation by Pseudomonas aeruginosa Phenazines. mBio 4, 1–9. 10.1128/mBio.00526-12.Editor

Norman, T.M., Lord, N.D., Paulsson, J., Losick, R., 2013. Memory and modularity in cell-fate decision making. Nature 503, 481–486. 10.1038/nature12804

Onaka, H., Mori, Y., Igarashi, Y., Furumai, T., 2011. Mycolic Acid-Containing Bacteria Induce Natural-Product Biosynthesis in Streptomyces Species. Appl. Environ. Microbiol. 77, 400–406. 10.1128/AEM.01337-10

Orazi, G., O’Toole, G.A., 2017. Pseudomonas aeruginosa Alters Staphylococcus aureus Sensitivity to Vancomycin in a Biofilm Model of Cystic Fibrosis Infection. mBio 8, e00873–17. 10.1128/mBio.00873-17

Piewngam, P., Zheng, Y., Nguyen, T.H., Dickey, S.W., Joo, H.S., Villaruz, A.E., Glose, K.A., Fisher, E.L., Hunt, R.L., Li, B., Chiou, J., Pharkjaksu, S., Khongthong, S., Cheung, G.Y.C., Kiratisin, P., Otto, M., 2018. Pathogen elimination by probiotic Bacillus via signalling interference. Nature 562, 532–537. 10.1038/s41586-018-0616-y

Pisithkul, T., Schroeder, J.W., Trujillo, E.A., Yeesin, P., Stevenson, D.M., Chaiamarit, T., Coon, J.J., Wang, J.D., Amador-Noguez, D., 2019. Metabolic Remodeling during Biofilm Development of Bacillus subtilis. mBio 10, e00623–19. 10.1128/mBio.00623-19

Popp, P.F., Dotzler, M., Radeck, J., Bartels, J., Mascher, T., 2017. The Bacillus BioBrick Box 2.0: Expanding the genetic toolbox for the standardized work with Bacillus subtilis. Sci. Rep. 7, 1–13. 10.1038/s41598-017-15107-z

Price, C.W., Fawcett, P., Cérémonie, H., Su, N., Murphy, C.K., Youngman, P., 2001. Genome-wide analysis of the general stress response in Bacillus subtilis. Mol. Microbiol. 41, 757– 774. 10.1046/j.1365-2958.2001.02534.x

Radeck, J., Kraft, K., Bartels, J., Cikovic, T., Dürr, F., Emenegger, J., Kelterborn, S., Sauer, C., Fritz, G., Gebhard, S., Mascher, T., 2013. The Bacillus BioBrick Box: Generation and evaluation of essential genetic building blocks for standardized work with Bacillus subtilis. J. Biol. Eng. 7. 10.1186/1754-1611-7-29

Ratnayake-Lecamwasam, M., Serror, P., Wong, K.-W., Sonenshein, A.L., 2001. *Bacillus subtilis* CodY represses early-stationary-phase genes by sensing GTP levels. Genes Dev. 15, 1093–1103. 10.1101/gad.874201

Rébora, K., Laloo, B., Daignan-Fornier, B., 2005. Revisiting Purine-Histidine Cross-Pathway Regulation in Saccharomyces cerevisiae: A Central Role for a Small Molecule. Genetics 170, 61–70. 10.1534/genetics.104.039396

Reilman, E., Mars, R.A.T., van Dijl, J.M., Denham, E.L., 2014. The multidrug ABC transporter BmrC/BmrD of Bacillus subtilis is regulated via a ribosome-mediated transcriptional attenuation mechanism. Nucleic Acids Res. 42, 11393–11407. 10.1093/nar/gku832

Richts, B., Rosenberg, J., Commichau, F.M., 2019. A Survey of Pyridoxal 5′-Phosphate-Dependent Proteins in the Gram-Positive Model Bacterium Bacillus subtilis. Front. Mol. Biosci. 6, 32. 10.3389/fmolb.2019.00032

Rosebrock, A.P., Caudy, A.A., 2017. Metabolite Extraction from Saccharomyces cerevisiae for Liquid Chromatography – Mass Spectrometry 1–6. 10.1101/pdb.prot089086

Rosenthal, A.Z., Qi, Y., Hormoz, S., Park, J., Li, S.H.-J., Elowitz, M.B., 2018. Metabolic interactions between dynamic bacterial subpopulations. eLife 7. 10.7554/eLife.33099

Saxild, H.H., Brunstedt, K., Nielsen, K.I., Jarmer, H., Nygaard, P., 2001. Definition of the Bacillus subtilisPurR Operator Using Genetic and Bioinformatic Tools and Expansion of the PurR Regulon with glyA, guaC,pbuG, xpt-pbuX, yqhZ-folD, and pbuO. J. Bacteriol. 183, 6175–6183. 10.1128/jb.183.21.6175-6183.2001

Schultz, A.C., Nygaard, P., Saxild, H.H., 2001. Functional analysis of 14 genes that constitute the purine catabolic pathway in Bacillus subtilis and evidence for a novel regulon controlled by the PucR transcription activator. J. Bacteriol. 183, 3293–302. 10.1128/JB.183.11.3293-3302.2001

Scott, M., Gunderson, C.W., Mateescu, E.M., Zhang, Z., Hwa, T., 2010. Interdependence of Cell Growth and Gene Expression: Origins and Consequences. Science 330, 1099–1102. 10.1126/science.1192588

Shafikhani, S.H., Mandic-Mulec, I., Strauch, M.A., Smith, I., Leighton, T., 2002. Postexponential regulation of sin operon expression in Bacillus subtilis. J. Bacteriol. 184, 564–571. 10.1128/JB.184.2.564-571.2002

Shank, E.A., Klepac-ceraj, V., Collado-torres, L., Powers, G.E., Losick, R., 2011. Interspecies interactions that result in Bacillus subtilis forming bio films are mediated mainly by members of its own genus. PNAS 108, E1236–E1243. 10.1073/pnas.1103630108

Shivers, R.P., Sonenshein, A.L., 2004. Activation of the Bacillus subtilis global regulator CodY by direct interaction with branched-chain amino acids: CodY activation by amino acids. Mol. Microbiol. 53, 599–611. 10.1111/j.1365-2958.2004.04135.x

Sonenshein, A.L., 2007. Control of key metabolic intersections in Bacillus subtilis. Nat. Rev. Microbiol. 5, 917–927. 10.1038/nrmicro1772

Sonenshein, A.L., 2005. CodY, a global regulator of stationary phase and virulence in Gram-positive bacteria. Curr. Opin. Microbiol. 8, 203–207. 10.1016/j.mib.2005.01.001

Stacy, A., Everett, J., Jorth, P., Trivedi, U., Rumbaugh, K.P., Whiteley, M., 2014. Bacterial fight-and-flight responses enhance virulence in a polymicrobial infection. Proc. Natl. Acad. Sci. 111, 7819–7824. 10.1073/pnas.1400586111

Stubbendieck, R.M., Straight, P.D., 2017. Linearmycins Activate a Two-Component Signaling System Involved in Bacterial Competition and Biofilm Morphology. J. Bacteriol. 199, e00186–17. 10.1128/JB.00186-17

Stubbendieck, R.M., Straight, P.D., 2015. Escape from Lethal Bacterial Competition through Coupled Activation of Antibiotic Resistance and a Mobilized Subpopulation. PLoS Genet. 11, e1005722. 10.1371/journal.pgen.1005722

Teichman, G., Cohen, D., Ganon, O., Dunsky, N., Shani, S., Gingold, H., Rechavi, O., 2023. RNAlysis: analyze your RNA sequencing data without writing a single line of code. BMC Biol. 21, 74. 10.1186/s12915-023-01574-6

Traxler, M.F., Seyedsayamdost, M.R., Clardy, J., Kolter, R., 2012. Interspecies modulation of bacterial development through iron competition and siderophore piracy. Mol. Microbiol. 86, 628–644. 10.1111/mmi.12008

Tsui, W.H.W., Yim, G., Wang, H.H., Mcclure, J.E., Surette, M.G., Davies, J., 2004. Dual Effects of MLS Antibiotics : Transcriptional Modulation and Interactions on the Ribosome. Chem. Biol. 11, 1307–1316. 10.1016/j

Turner, R.J., Bonner, E.R., Grabner, G.K., Switzer, R.L., 1998. Purification and Characterization of Bacillus subtilis PyrR, a Bifunctional pyr mRNA-binding Attenuation Protein/Uracil Phosphoribosyltransferase *. J. Biol. Chem. 273, 5932–5938. 10.1074/jbc.273.10.5932

Turner, R.J., Lu, Y., Switzer, R.L., 1994. Regulation of the Bacillus subtilis pyrimidine biosynthetic (pyr) gene cluster by an autogenous transcriptional attenuation mechanism. J. Bacteriol. 176, 3708–3722. 10.1128/jb.176.12.3708-3722.1994

van Gestel, J., Vlamakis, H., Kolter, R., 2015. From cell differentiation to cell collectives: Bacillus subtilis uses division of labor to migrate. PLoS Biol. 13, e1002141. 10.1371/journal.pbio.1002141

Weng, M., Nagy, P.L., Zalkin, H., 1995. Identification of the Bacillus subtilis pur operon repressor. Proc. Natl. Acad. Sci. U. S. A. 92, 7455–7459. 10.1073/pnas.92.16.7455

Winstedt, L., von Wachenfeldt, C., 2000. Terminal Oxidases of Bacillus subtilisStrain 168: One Quinol Oxidase, Cytochromeaa3 or Cytochrome bd, Is Required for Aerobic Growth. J. Bacteriol. 182, 6557–6564. 10.1128/jb.182.23.6557-6564.2000

Wolfe, A.J., 2005. The acetate switch. Microbiol. Mol. Biol. Rev. MMBR 69, 12–50. 10.1128/MMBR.69.1.12-50.2005

Yang, J., Barra, J.T., Fung, D.K., Wang, J.D., 2022. Bacillus subtilis produces (p)ppGpp in response to the bacteriostatic antibiotic chloramphenicol to prevent its potential bactericidal effect. mLife 1, 101–113. 10.1002/mlf2.12031

Yannarell, S.M., Beaudoin, E.S., Talley, H.S., Schoenborn, A.A., Orr, G., Anderton, C.R., Chrisler, W.B., Shank, E.A., 2023. Extensive cellular multi-tasking within Bacillus subtilis biofilms. mSystems 8, e00891–22. 10.1128/msystems.00891-22

Yasbin, R.E., Young, F.E., 1974. Transduction in Bacillus subtilis by bacteriophage SPP1. J. Virol. 14, 1343–1348. 10.1128/JVI.14.6.1343-1348.1974

Zhu, B., Stülke, J., 2018. SubtiWiki in 2018: From genes and proteins to functional network annotation of the model organism Bacillus subtilis. Nucleic Acids Res. 46, D743–D748. 10.1093/nar/gkx908

